# The CryoEM Structure of Human GPR75: Insights into ECL2-Mediated Activation

**DOI:** 10.1101/2025.07.30.667585

**Authors:** A. Manuel Liaci, Vasudha Gathey, Aniket Magarkar, Tobias Kiechle, Camilla Mayer, Giuseppe Bruschetta, Jürgen Schymeinsky, Curtis R. Warren, Herbert Nar, Rebecca Ebenhoch

## Abstract

GPR75 is one of the most promising emerging targets for the treatment of obesity and related co-morbidities, as its loss of function directly correlates with a decreased body mass index (BMI) in humans. To date, little is known about the structure and underlying biology of GPR75, which is classified as an orphan GPCR and has low homology to other GPCRs.

Here, we describe the cryoEM structure of ICL3-BRIL-fused, unliganded human GPR75, together with functional data on G-protein coupling and effects of its putative ligands. GPR75 is present in an active-like state probably induced by the structure of its extracellular loop 2 (ECL2) which folds back deeply into the orthosteric pocket. This feature is important for both receptor integrity and signaling, as shown by mutagenesis studies. This structural finding is consistent with moderate constitutive activity for Gα_i1_, while Gα_q_ does not seem to be recruited. Furthermore, functional data indicate that the putative ligands 20-HETE and CCL5 have no measurable effect on GPR75.

**Significance Statement:** Obesity is a major global health crisis which will affect a quarter of the world’s population by 2035. Many existing therapies have significant shortcomings, including emetic effects and the need for frequent injections. GPR75 is an understudied orphan GPCR which has recently emerged as one of the most promising novel obesity targets based on large-scale human genetic studies. However, the sparse knowledge we currently have about this receptor presents a major roadblock for the development of effective GPR75-based therapies. This study investigates the basic biology of GPR75 on a molecular level, including its structure and downstream signaling. These findings provide the groundwork for future efforts targeting GPR75 to fight the obesity pandemic.

## Introduction

GPR75 is a G-protein coupled receptor (GPCR) expressed in the brain, particularly in POMC neurons and astrocytes in hypothalamic regions such as the arcuate nucleus (1–6). Despite being attributed to GPCR class A, GPR75 has a unique sequence that bears little resemblance to any other GPCR. As such, GPR75 shows a low conservation of most canonical microswitches and amino acid motifs (7), as well as an unusually long disordered intracellular loop 3 (ICL3) and C-terminus.

GPR75 has emerged as an exciting new target in obesity, as an exome-sequencing study of 645,626 individuals found a genetic link between GPR75 loss of function and reduced body mass index (BMI) in humans. GPR75 knockout mice are protected from diet-induced weight gain, suggesting a fundamental role in the regulation of body weight (2). The observed BMI reduction has been attributed to a reduced food intake (3, 4, 8) and effects on energy expenditure (9), and correlates with improved insulin sensitivity (2, 8, 9). Unlike most other genes that promote leanness upon disruption, GPR75 loss of function also protects mice from developing non-alcoholic fatty liver disease (4, 10). Apart from its role in obesity, GPR75 is highly expressed in the retina and may be involved in the development of age-related macular degeneration (11). Additionally, GPR75 expressed in the lung might contribute to the pathogenesis of pulmonary hypertension (12).

Links to other morbidities and inflammatory processes have largely been extrapolated from the functions of GPR75’s putative ligands (9, 12–18). Two structurally diverse physiological ligands for GPR75 have been proposed in the literature: the eicosanoid 20-hydroxyeicosatetraenoic acid (20-HETE), and the cytokine CC-motif chemokine 5 (CCL5, RANTES) (15, 17–20). However, the available data on these ligands do not fulfill the IUPHAR criteria for GPCR deorphanization (21), and the underlying biology of GPR75 remains poorly understood. According to these studies, GPR75 signals through the G-protein Gα_q_ (15, 17–20), whereas an independent study detected constitutive signaling through Gα_i1_, but not Gα_q_ (22).

Here, we present the structure of unliganded (apo) GPR75 solved by cryo-electron microscopy (cryoEM). The apo GPR75 is present in what we define as an ‘active-like’ state. Neither 20-HETE nor CCL5 show any effect on GPR75, but we observed moderate constitutive activity mediated through Gα_i1_. The orthosteric site of GPR75 is prominently filled with its own backfolded ECL2, which is crucial to GPR75‘s structural integrity and promotes constitutive signaling.

## Results

### Description of the GPR75 apo structure

#### Overall structure

We solved the structure of human GPR75 in absence of a ligand or G-protein using single particle cryoEM, with the local resolution of the receptor at about 3 Å (**Figure 1** & **Figure S1**, **Table S1**). A fragment antigen-binding (Fab) directed against the apocytochrome b562 (BRIL)-insert in intracellular loop 3 (ICL3) and an anti-BRIL nanobody (Nb) were used to facilitate particle alignment (**Figure 1A,B** & **S1A,B**) (23, 24).

**Figure 1.**
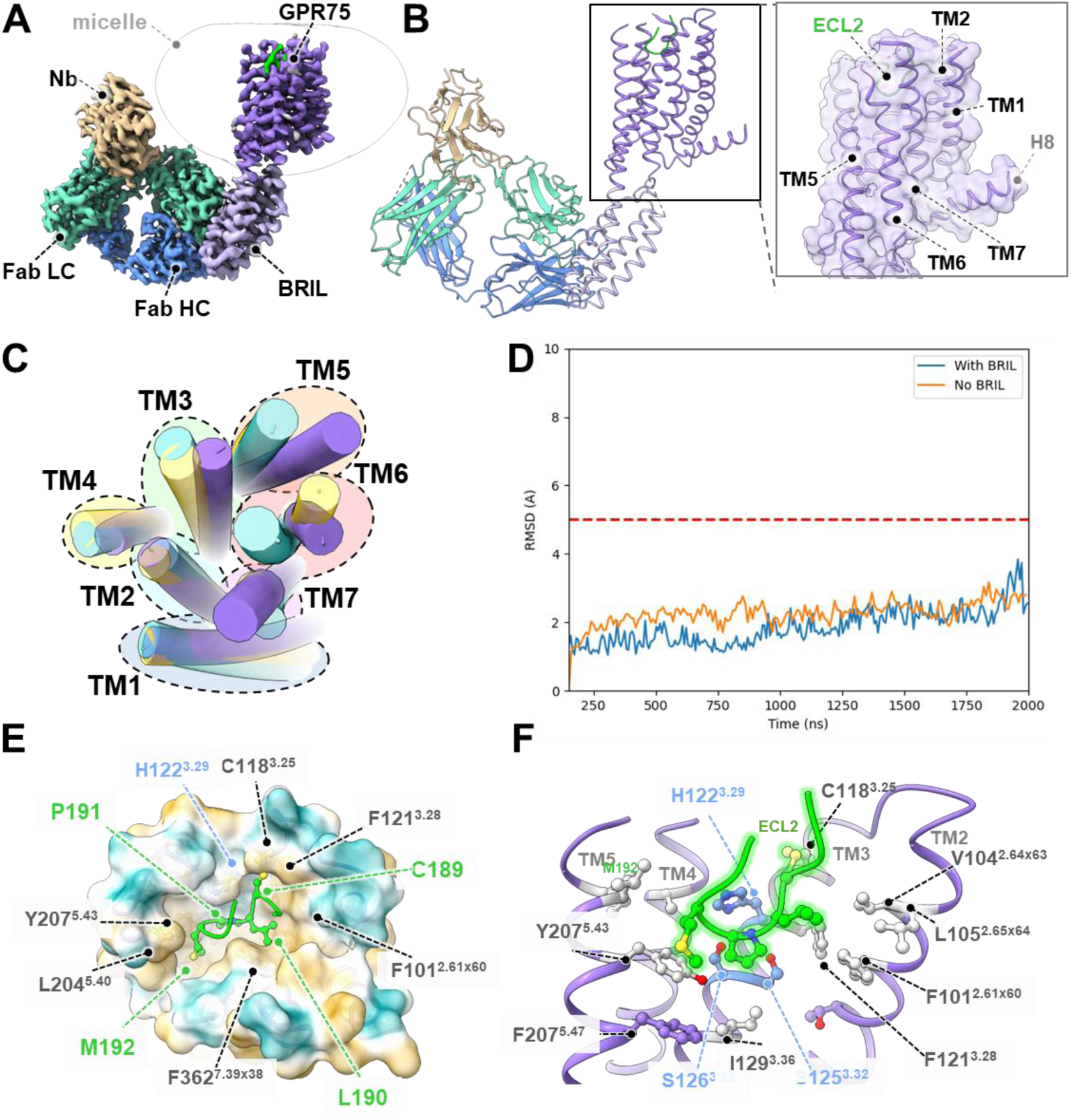
CryoEM structure of human GPR75. **A** Composite cryoEM map of the GPR75-BRIL : anti-BRIL-Fab : anti-Fab-Nb complex. The resolution allows for confident placement of side chains. **B** Atomic model of the complex, colored as in A. Zoom-in shows a detailed view of the TM helix arrangement. **C** Comparison of the GPR75 (purple) tertiary structure to the active (PDB 7BU7, yellow) and inactive (7BVQ, teal) states of β1-AR. Depicted are the G-protein binding cavities of GPR75 and β1-AR as seen from the intracellular side. H8 and intracellular loops were omitted for clarity. **D** MD simulation of GPR75 in a lipid bilayer with an ICL3-BRIL insertion (blue) vs wild-type ICL3 (orange), starting from the experimental conformation. **E** Top view of the orthosteric pocket. The pocket is mostly hydrophobic and filled with four residues of ECL2. The surface is colored by hydrophobicity from gold (20) to teal (−20). **F** Side view of the orthosteric pocket. TM5 & TM6 are omitted for clarity.

Class A GPCRs are thought to possess a common cascade of structural rearrangements during activation involving 34 amino acids in conserved motives such as CWxP, PIF, the sodium binding pocket, NPxxY, and DRY (7). However, GPR75 and its most closely related class A GPCRs harbor merely 10-13% global sequence identity and 23-24% sequence similarity and display poor conservation of these motifs and microswitch residues (25). We used a machine learning (ML) approach that annotates the conformation of class A GPCRs based on a training set of thousands of conformations sampled from 38 unique class A structures to tentatively characterize the experimentally determined conformation of apo GPR75 (26). To our surprise, the algorithm reported a 98% probability that the observed conformation of unliganded GPR75 is active-like. One of the most closely related GPCRs is human β1-adrenoceptor (β1-AR) with only 12% sequence identity and 23% sequence similarity. We nonetheless chose β1-AR for in-depth structural comparisons and visualizations of the conformational state of GPR75, as it features a reasonable sequence similarity in the functionally most important amino acid positions (25) and structural information of its various activity states is available (27). When aligned with the active and inactive states of β1-AR, the apo GPR75 conformation shows hallmarks of the active state of β1-AR, with a swung-out TM6 and extensive contacts between TM3 and TM7 (**Figure 1C, Figure S2A-C**). Accordingly, the structural agreement between apo GPR75 and active β1-AR (TM score 0.81 / rmsd 3.2 Å) is higher than for inactive β1-AR (0.76 / 3.8 Å).

There is a possibility that the BRIL insertion into ICL3 might coax GPR75 into the observed conformation. However, GPCRs solved by cryoEM with rigid BRIL-fusions have so far only been observed in inactive and intermediate conformations (28–32), in line with the energy landscape theory put forth by Kobilka and colleagues (33–35). In order to further probe the resting conformation of GPR75, we performed molecular dynamics (MD) simulations in a simulated lipid bilayer over 2 us. To this end, we compared the conformational space of GPR75 with and without the BRIL fusion by tracing the rmsd starting from our observed conformation (**Figure 1 D**). According to the simulation, GPR75 stably retains its conformation over 2 µs regardless of the presence or absence of the BRIL insertion. In addition, other authors find that a different approach inserting a non-rigid BRIL at a more distal position in GPR75’s TM5 & TM6 results in a highly similar state (36). The authors also raised a nanobody against recombinant GPR75 using yeast display, which bound to the observed conformation. This finding was unexpected, as nanobodies raised by display methods tend to sample the predominant conformations of the target molecule. Taken together, the fact that GPR75 can stably retain this conformation in absence of the BRIL fusion and that similar conformations can be obtained with different stabilization approaches indicates that the conformation we observe is not induced by the BRIL insertion, although the insertion does increase protein expression and *in vitro* longevity.

#### Conserved motifs and microswitches

Complementing the overall structure, the conservation and conformation of important motives and microswitches is a critical factor in assessing the activity state of a class A GPCR. The resolution and quality of our cryoEM map is sufficient to allow us to analyze these motives in detail. Residue Y207^5.43^ is stacked against M192^ECL2^ of the CLPM motif and seals off the bottom of the orthosteric pocket (superscripts refer to the Ballesteros-Weinstein nomenclature (37)) (**Figure 1E-F**). Underneath this residue, bridged by F211^5.47^, are the locations of the CWxP and PIF motifs. In GPR75, however, these motives are poorly conserved. The only nominally and functionally conserved residue of the CWxP motif is P336^6.50^ (P365 in our BRIL-fusion construct), which induces a kink in TM6 consistent with its canonical function, promoting the observed flexibility of this helix (**Figure 2, Figure S2I**). Position 6.48 is canonically occupied by a tryptophan and thought to be the ‘toggle switch’, one of the most studied microswitch residues in GPCRs (38). However, the residue is not conserved in GPR75 and instead replaced by the smaller C334^6.48^. The rotation of TM6 at this position is comparable to the active state of β1-AR (**Figure 2A-B**). The space into which the toggle switch would normally extend is occupied by TM7 residue I365^7.45^, which forms direct hydrophobic contacts with TM3 residue I129^3.36^ and thus contributes to the extensive TM3-TM7 interface. Hence, even the sterically small sidechain of C335^6.48^ might have less freedom to rotate into the ‘inactive’ conformation.

**Figure 2.**
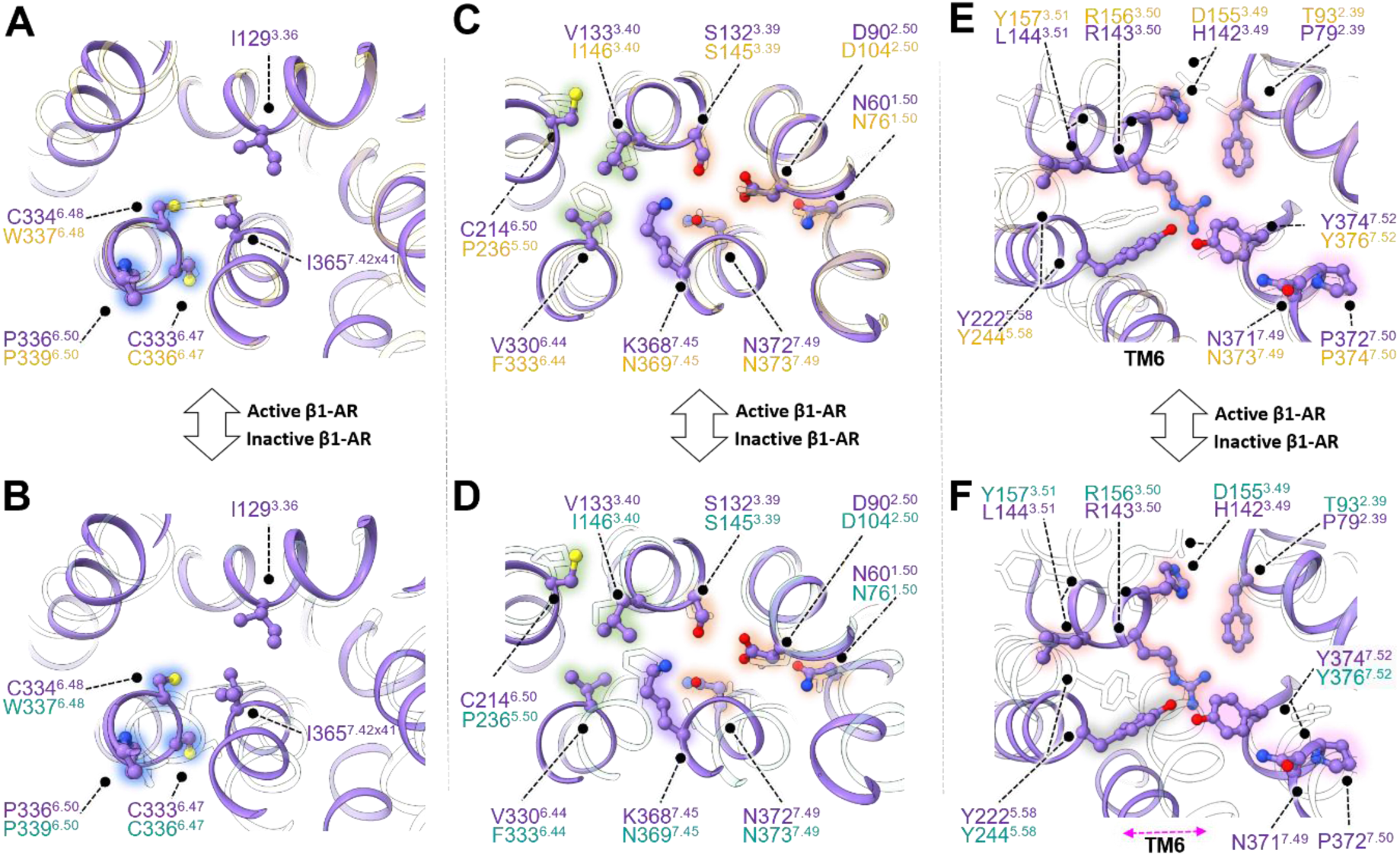
Structural motifs and microswitches of apo GPR75 compared to the active (top row) and inactive (bottom row) states of β1-AR. The superposed active (PDB 7BU7) and inactive (7BVQ) state structures of β1-AR are shown as white outlines. Important amino acids are shown as sticks and labeled (purple – GPR75; white/yellow – β1-AR active; white/teal – β1-AR inactive). View is from the extracellular side. **A,B** The CWxP motif is poorly conserved in GPR75 and rotated as in the active state of β1-AR, allowing additional interactions between TM7 and TM3. **C,D** The sodium binding pocket of GPR75 (orange highlight) is collapsed and features a non-canonical lysine residue at position 368^7.45^ (purple highlight). The PIF motif (green highlight) is poorly conserved. **E,F** Residues R143^3.50^, Y222^5.58^, and Y376^7.53^ form an interacting network that is typical for active GPCR states. Residues of the NPxxY (pink highlight) and DRY motif (red highlight) as well as a swung-out TM6 (pink arrow) bear hallmarks of an active GPCR.

Much like the CWxP motif, the PIF motif is also not conserved in GPR75 (**Figure 2C-D**). This motif is thought to initiate the structural rearrangements leading to receptor (in)activation. In particular, the canonical phenylalanine is replaced by C214^5.50^, which is only present in four otherwise unrelated class A GPCRs (LGR5-6, olfactory receptor 1G1, and MRGPRD). Close to the non-conserved PIF motif is the location of the sodium binding pocket (**Figure 2C-D**). This set of residues appears to be collapsed, which is a hallmark of activated GPCRs, and there is neither space nor observed density for a sodium ion. A particularly striking feature of GPR75 is the presence of a large, positively charged lysine (K368) at position 7.45 whose amine group lies in the center of the motif. The presence of this residue raises the question if the GPR75 sodium binding pocket is at all capable of adapting an ‘inactive’ configuration that canonically binds sodium ions. Mutating this residue to an alanine completely abrogated protein expression (data not shown).

Lastly, the DRY motif canonically stabilizes a GPCR in its inactive state, particularly through the ‘ionic lock’, an interhelical salt bridge between positions 3.50 and 6.30. While H142^3.49^ and L144^3.51^ are non-canonical in GPR75, the ionic lock pair of R143^3.50^ and D316^6.30^ (D345 in our structure) is in fact conserved. In the structure, D316^6.30^ is not well-resolved due to high flexibility, but its location is far away from R143^3.50^, similar to the active state of β1-AR. Instead, R143^3.50^ extends toward the membrane and forms an interaction network with Y222^5.58^ and Y376^7.53^ from the conserved NPxxY motif (**Figure 2E-F**). However, it is possible that the local conformation of D316^6.30^ and its neighboring residues is influenced by the proximity to the BRIL insertion.

In summary, we define the observed conformation of GPR75 as ‘active-like’ based on the comparison to other GPCRs and structural data accumulated by us and others (36). However, in light of the unique sequence of GPR75, it might be reasonable to question the extent to which the rules delineated for class A activation are at all relevant in this case. GPR75’s overall structure, microswitch architecture, and particularly its molecular dynamics suggest that the conformation we observe is the resting state of GPR75 and that this conformation is driven by the GPCR rather than the fusion protein.

#### Detailed description of the orthosteric site

We were able to model the side chains of the orthosteric pocket (**Figure 1E-F**) excepting the out-facing C-terminal portion of TM6, which is poorly ordered starting at P365 in our BRIL-fusion construct (corresponding to P336^6.50^ in the wildtype (wt) sequence). The orthosteric pocket is mostly hydrophobic, and in particular the bottom of the cavity is lined by exclusively hydrophobic residues. A hydrophilic patch formed by the TM3 residues H122^3.29^, S125^3.32^, and S126^3.33^ is located on the lateral wall (**Figure 1E, Figure S2D**).

Perhaps the most striking feature of the structure is GPR75’s ECL2, which is folded back deeply into the orthosteric site (**Figure 1E-F, Figure S2E-F**). The loop is kept in place by a canonical disulfide bridge between residues C118^3.25^ and C189^ECL2^. While the hydrophobic side chains L190^ECL2^, P191^ECL2^, and M192^ECL2^ make extensive contacts with the hydrophobic bottom of the orthosteric site, their backbone is stabilized by the polar patch of TM3 residues, particularly H122^3.29^. Together, these four ECL2 residues form a stretch we termed the ‘CLPM motif’, whose ‘CLP’ residues are highly conserved among species (**Figure S2G, S5**). The side chain M192^ECL2^ extends into a hydrophobic sub-pocket (**Figure 1E**), but the residue is partially open to the membrane and extracellular space due to the apparent flexibility of TM6. The residue appears to possess some degree of freedom, as the cryoEM map shows an additional density protruding from the C_α_ of M192^ECL2^ which might represent an alternative conformation (**Figure S2H**). While the ‘CLP’ residues are strictly conserved, other species also feature other bulky hydrophobic sidechains (leucine, valine, isoleucine) or histidine at position 192^ECL2^ (**Figure S5**).

We reasoned that, in concoction with the ‘active-like’ conformation of GPR75, the ECL2 self-engagement might indicate increased constitutive activity, which led us to investigate the GPCR’s constitutive activity profile with respect to the G-protein binding partners reported in the literature.

### GPR75 displays moderate constitutive activity for Gα_i1_

The G-protein coupling of GPR75 remains controversial, as several publications report a ligand-induced coupling to Gα_q_ (15, 17–20, 39), while two recent publications suggest constitutive signaling through Gα_i1_ (4, 22). For an initial assessment, we compared our structure to a modeled complex of GPR75 with the trimeric G-protein consisting of the three subunits Gα_i1_, Gβ_1_, and Gγ_2_ (Gα_i1_β_1_γ_2_) prepared with ColabFold (40) (**Figure S2J-L**). Except for the extracellular loops, the computed active state overlaps remarkably well with the experimentally determined coordinates, and upon superposition the Gα_i1_ C-terminus can be accommodated without major steric clashes.

Prompted by the active-like state observed in our unliganded structure, we tested the constitutive activity of GPR75 for these two G-proteins using a BRET2 assay (41–45) (**Figure 3**). S1P3, the positive control for Gα_q_ coupling (46) showed a clear reduction in BRET2 signal compared to the mock control transfected without GPCR, indicating constitutive receptor activity (**Figure 3A**). In contrast, GPR75 and the negative control GPR52 (22) exhibited no such effect. However, GPR75 showed a significant effect on Gα_i1_ coupling, although the effect is rather low compared to the positive control GPR20 (47) (**Figure 3B**). We confirmed the observed low constitutive Gα_i1_ signaling in a cyclic AMP (cAMP) assay in CHO-K1 cells. We have been able to consistently reproduce data showing a 1.5- to threefold change in the IC_50_ values between parental and GPR75-expressing cells, indicating inherently lower cAMP levels in the GPR75-overexpressing cells (**Figure 3C, S3A-B**). Thus, our data suggest that GPR75 possesses low constitutive activity for Gα_i1_ but not Gα_q_. We have so far been unable to produce a stable complex between GPR75 and Gα_i1_ *in vitro*, which might indicate a low coupling affinity overall.

**Figure 3.**
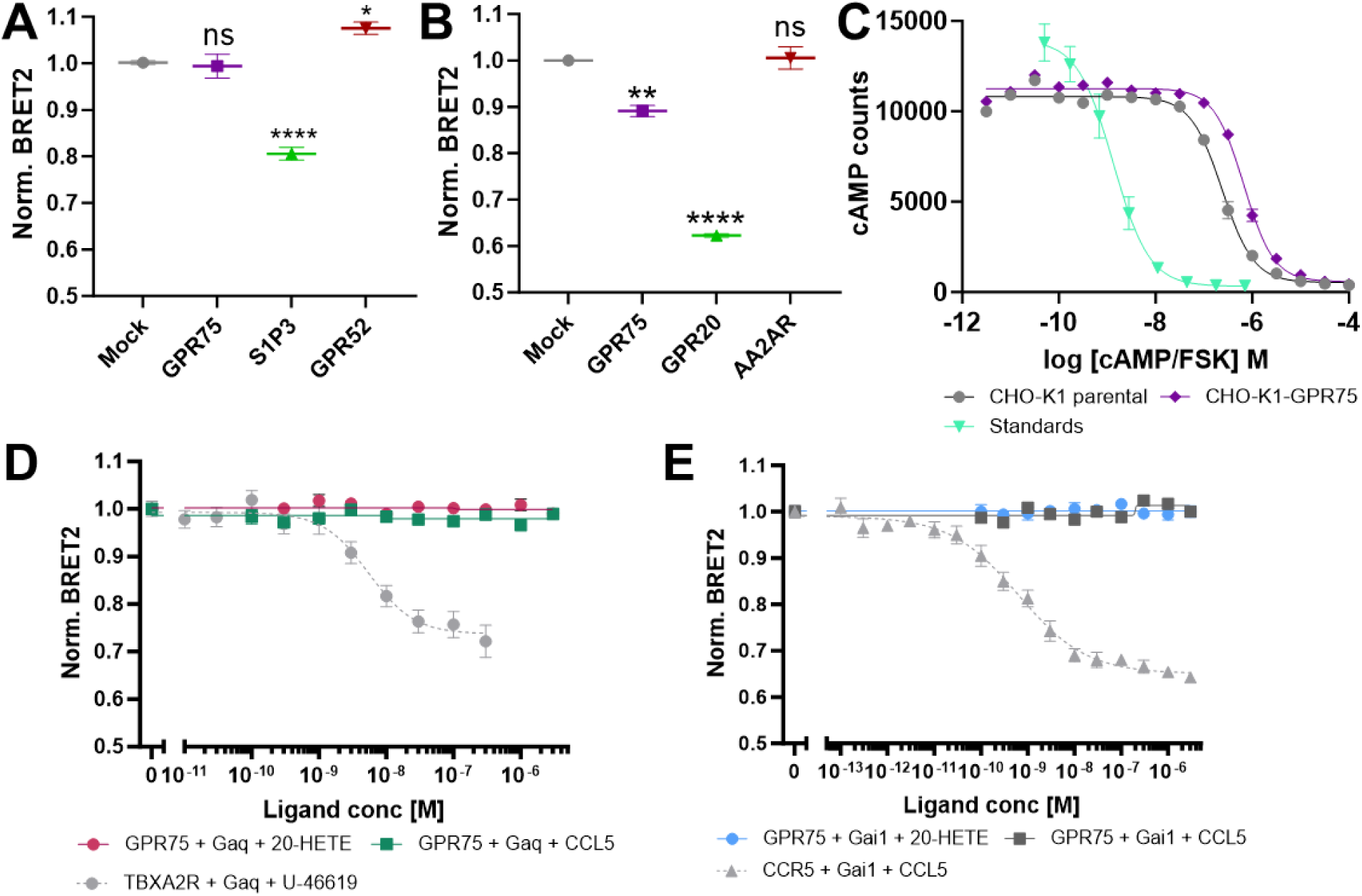
Constitutive and ligand-induced activation of GPR75. **A** BRET2 assay examining the constitutive activity of GPR75 for Gα_q_. S1P3 (46) and GPR52 (22) were included as positive and negative controls for Gα_q_ coupling, respectively. **B** BRET2 assay examining the constitutive activity of GPR75 for Gα_i1_. GPR20 and AA2AR were included as positive and negative controls, respectively (47). **A-B** Normalized BRET2 values ±SEM from three or more biologically independent experiments with six technical replicates each. Statistical difference to Mock was tested using one-way ANOVA followed by Dunnett’s *post-hoc* test (****P<0.0001, **P<0.01, *P<0.05, n.s.>0.05). **C** cAMP assay examining the constitutive activity of GPR75 for Gα_i1_. GPR75 over-expression leads to decreased cAMP levels in CHO-K1 cells. Representative figure from an experiment performed with 750 cells/well. **D** BRET2 assay examining the ligand-induced activation of GPR75 for Gα_q_. TBXA2R and its antagonist U-46619 were included as technical control (EC50: 5.46×10^-9^M). **E** BRET2 assay examining the ligand-induced activation of GPR75 for Gα_i1_. The CCR5-CCL5 pair was included as a technical control. **D-E** Normalized BRET2 values ±SEM from three or more biologically independent experiments with three technical replicates each.

### 20-HETE and CCL5 have no effect on GPR75 G-protein signaling

In addition to profiling the constitutive activity of GPR75 for Gα_i1_ and Gα_q_, we aimed to reconfirm the effect of GPR75’s endogenous ligands. Two chemically very different endogenous agonists have been reported for GPR75, both reportedly signaling through Gα_q_ with picomolar K_D_: The lipidic eicosanoid 20-HETE (20, 39) and the proteinaceous chemokine CCL5 (17, 18).

To further assess the functionality of these molecules as GPR75 agonists, we performed a BRET2 assay in agonist mode. The integrity and activity of both 20-HETE and CCL5 were confirmed in orthogonal quality control assays (**Figure S3C-F**, **S6-7)**. However, neither ligand had any effect on GPR75 signaling through Gα_q_ even at micromolar concentrations (**Figure 3D**). As an orthogonal confirmatory measure, we also performed a Calcium flux assay in which neither molecule elicited a cellular response at concentrations up to the high nanomolar range (**Figure S3G**). Lastly, we also tested the possibility of 20-HETE- or CCL5-induced signaling through Gα_i1_ in a BRET2 assay, which also showed no effect (**Figure 3E**). An independent confirmation of these results was conducted by monitoring the direct effect of both ligands on the thermostability of GPR75-BRIL as a measure of direct target engagement (**Figure S3H-I**). However, neither the addition of 9 μM 20-HETE nor 6 μM CCL5 resulted in a noticeable change in the GPR75 melting point.

To further assess how these two ligands might bind to GPR75, we compared the architecture of the GPR75 orthosteric site to that of lipid GPCRs such as the putative micromolar-affinity 20-HETE receptor FFAR1 (GPR40) (48) (**Figure S3J-L**). Lipid-binding GPCRs generally feature an ECL2 cap that blocks access by soluble ligands, positively charged patches able to form salt bridges with the lipid’s terminal carboxyl groups, and access to the orthosteric site through lateral gaps between their TM1/7 or TM4/5 helix pairs. While GPR75 possesses a hydrophobic orthosteric site that is shielded from the exterior by ECL2 and a small polar patch formed by H122, a potential lipid binding would likely have to be accommodated in a novel fashion. As such, the mobile TM6 might allow passage through a gap formed with TM5, and the polar patch formed by H122 might counteract the charge of carboxylic headgroups in the ligands, although it is unknown to which extent this residue can be protonated. Lastly, the GPR75 ECL2 shields the pocket from the extracellular space – however, the tight packing and deep burial of the CLPM motif would certainly pose a considerable obstacle for ligands to overcome.

CCL5 binds a set of different chemokine receptors (CCR1,3,4,5) and has the highest affinity for CCR5 (49). We therefore compared GPR75 to the structure of CCL5 bound to CCR5 (50) (**Figure S3M-N**). In the structure, CCL5’s extended N-terminus stretches into an open extracellular vestibule formed by an outward movement of the CCR5 TM2. This TM2 movement is a hallmark of the chemokine-receptor interactions (51), as shown for other GPCRs such as CXCR4, CXCR2, and US28 (52–54). Typical for peptide-binding GPCRs, the ECL2 of CCR5 does not obstruct the orthosteric site and instead forms a hairpin (55), and the N-terminus of CCL5 accommodates a large tyrosine residue into the space opened by TM2. The orthosteric pocket of GPR75 bears little resemblance to that of CCR5 in terms of sequence, charge, and topology, and an analogous CCL5 binding mode would only be possible if the chemokine were to induce major conformational changes in GPR75, displacing both its TM2 and ECL2 – a scenario which appears unlikely given the tight binding of ECL2. A fitting example for this scenario is BILF1, which directly descended from the chemokine receptors but lost the ability to bind chemokines due to its self-binding ECL2 (56).

In summary, both 20-HETE and CCL5 do not act as GPR75 ligands for G-protein signaling in our functional assays. The structural data imply that the binding sites of known receptors for these two molecules are not functionally conserved in GPR75, further indicating that it is unable to functionally interact with either candidate ligand.

### The GPR75 ECL2 is important for receptor integrity and conformation

One of the most intriguing features of the GPR75 apo structure is the backfolded CLPM motif in ECL2 that completely occupies the orthosteric site. Based on the well-ordered density of our structure, ECL2-backfolding is likely permanent in its unliganded state. In combination with the absence of a validated natural ligand and the observed active-like state of the apo GPR75, the prominent presence and high conservation of this motif raise several interesting questions as to its purpose. Three (not mutually exclusive) scenarios are conceivable that could explain its function: (I) ECL2 may form a tight lid above the otherwise very hydrophobic orthosteric pocket, modulating the response to (potentially unknown) ligands and preventing unspecific binding of lipophilic molecules from the extracellular space, (II) ECL2 backfolding might be a structural component of receptor stability and integrity, and (III) ECL2 might lead to a self-activation of GPR75.

As a first experiment, we performed MD simulations of GPR75 over the course of 2 µs, starting from our observed conformation with either the observed, deeply embedded ECL2 conformation or a starting model where ECL2 has been artificially ejected (**Figure S4A-B**). As evident from the data, artificial ECL2 ejection leads to a conformational destabilization, indicating that it contributes to the conformational uniformity of the active-like resting state. To further narrow down the options, we tested the effect of mutations in the CLPM motif on GPR75 expression, stability, and the constitutive activation of Gα_i1_ (**Figure 4**). To this end, we overexpressed a set of BRIL-fused point mutants and studied their *in vitro* thermostability in a nanoDSF experiment (**Figure 4A-B, Figure S4C**). Indeed, all mutations except M192A showed a decrease in thermostability or even a complete loss of melting transition, indicating a severe receptor destabilization. In addition, the protein yields and monodispersity upon overexpression were impaired, indicating reduced protein surface levels (**Figure S4C-E**). The strongest effect on both expression yields and thermostability were observed when mutating C198^ECL2^ and P191^ECL2^. These findings can be rationalized, as the loss of the canonical disulfide bridge between ECL2 and TM3 is known to destabilize GPCRs, and the mutation of P191^ECL2^ likely distorts the backbone conformations that are necessary for ECL2 to fully reach into the orthosteric site. Mutating the complete CLPM motif also had a severely disruptive effect on receptor expression and thermostability (**Figure 4A-B, Figure S4C-E**). We were able to recapitulate these findings in an MD simulation with both a P191A and a CLPM->CAAA mutation, both of which showed rapid loss of the resting state conformation (**Figure S4F-G**). In contrast, mutating the two residues L190^ECL2^ and M192^ECL2^ to alanine likely does not lead to a severely altered ECL2 conformation and hence displays a less pronounced destabilizing effect. We observe a possible alterative conformation for the side chain of M192^ECL2^ in our cryoEM map, indicating that this side chain does not contribute crucial interactions (**Figure S2H**).

**Figure 4.**
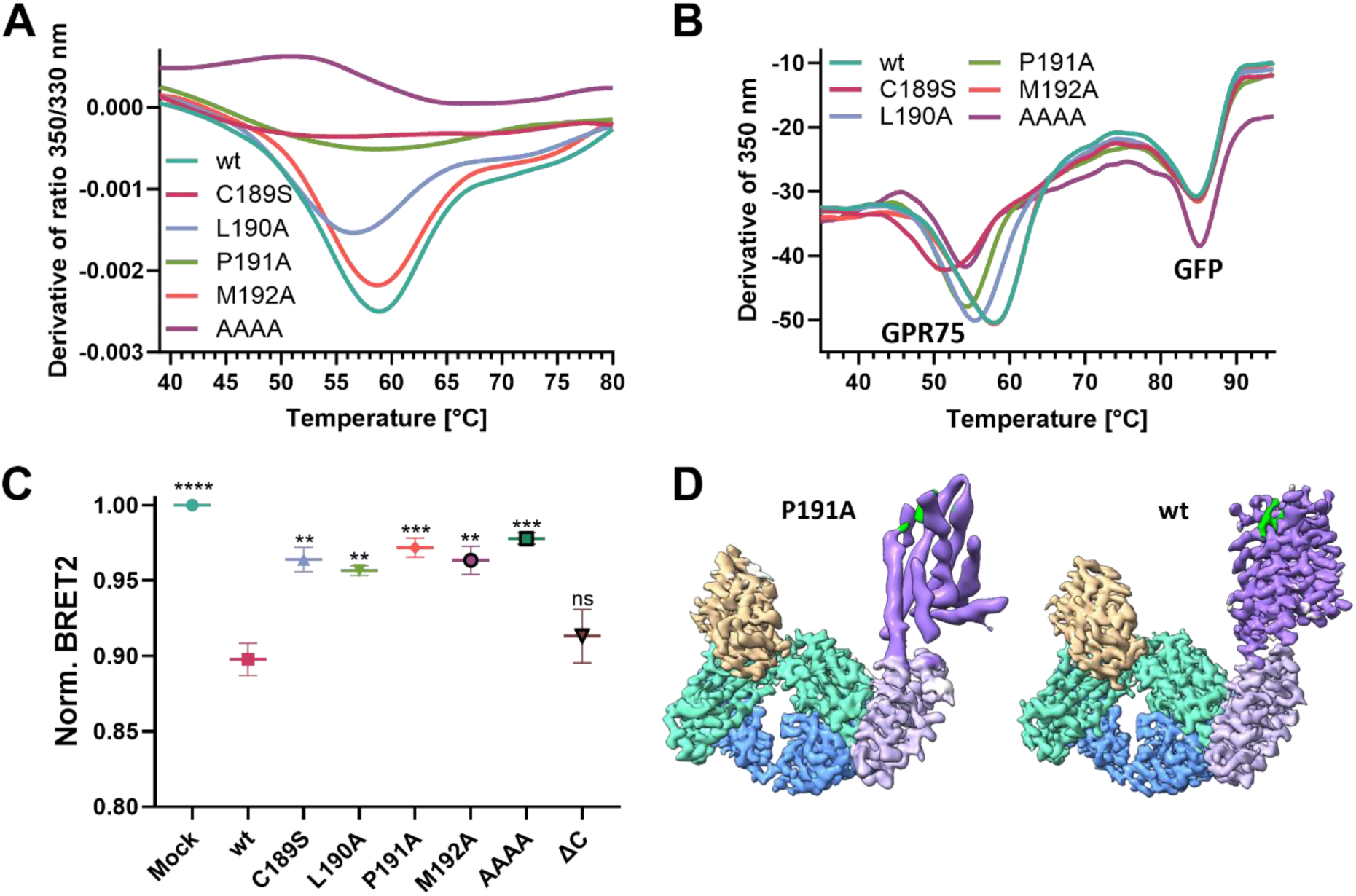
Effects of GPR75 ECL2 mutations. **A**,**B** The thermostability of BRIL-fused ECL2 mutations was assessed by nanoDSF. **C** BRET2 assay examining the constitutive activity of GPR75 ECL2 mutants for Gα_i1_. Normalized BRET2 values ±SEM from three or more biologically independent experiments with six technical replicates each. Statistical difference to GPR75 wt was tested using one-way ANOVA followed by Dunnett’s *post-hoc* test (****P<0.0001, ***P<0.001, **P<0.01, ns>0.05). **D** The structure of the GPR75 P191A mutant (left, colored as in Fig. 1) shows a poorly ordered GPCR (purple density) compared to the wt (right), indicating increased conformational flexibility. Both maps are rendered at a surface level of 0.29.

Lastly, we explored the functional consequence of ECL2 mutations in a BRET2 assay (**Figure 4C**). In the absence of a verifiable ligand, we assessed their effect on the constitutive activation of Gα_i1_. Indeed, all mutations result in a statistically significant reduction of constitutive activity for Gα_i1_ when compared to the wt, although we did not observe a complete activity loss. Since it is difficult to deconvolute to what extent this effect is caused by the reduced surface availability and stability of the GPR75 mutants, we collected cryoEM data of the unliganded P191A mutant to directly assess the effect of the mutation on the GPCR’s conformation (**Figure 4D**). Although expression produced lower yields than wt GPR75 (**Figure S4D-E**), the sample was stable enough to allow structure determination. In line with the MD data, the P191A mutation appears to affect the conformational uniformity of the GPCR, which now appears able to adopt a non-discrete range of conformations. As a result, focused refinement on the GPCR carried out in analogy to GPR75 wt does not reach near-atomic resolution, although the Fab-Nb portion is well resolved (**Figure 4D**). Based on the tertiary arrangement of the helices, the expanded conformational space appears to oscillate around the active-like state rather than snapping into an inactive state (**Figure S4H**). There is remaining observable density for ECL2 in the orthosteric site, indicating that the mutation does not result in a complete ejection of ECL2 from the pocket.

Taken together, these results indicate that the backfolding of ECL2 into the GPR75 orthosteric pocket is a feature that increases the receptor’s stability and expression levels, and also leads to increased conformational uniformity. Mutating ECL2 negatively affects the constitutive activity of GPR75, likely due to a combined effect of the loss of functional protein on the cell surface and on the increased conformational heterogeneity of the active-like state.

## Discussion

### GPR75 conformation

This work presents the high-resolution structure of the orphan GPCR GPR75 in its unliganded (apo) state. Both the global and site-specific features of the structure imply that apo GPR75 is present in what we define as an ‘active-like’ conformation: A swung-out and rotated TM6 that lacks the canonical W^4.48^ microswitch, extensive contacts between TM3 and TM7, a collapsed and potentially dysfunctional sodium binding pocket containing a rare, positively charged lysine residue, a broken ionic lock and a stacking interaction of R143^3.50^, Y222^5.58^, and Y376^7.53^, as well as a G-protein binding cavity theoretically fit to harbor a G-protein without major clashes. In line with this analysis, STAGS, an ML-based approach for the characterization of class A GPCR conformations (26), reports a 98% probability that unliganded GPR75 adopts an active-like conformation. In particular, the set of non-canonical alterations to the normally conserved motifs seem to promote the active-like state. However, in absence of a full agonist or antagonist, and without structurally characterized homologous GPCRs available, we lack the tools to experimentally assess the conformational boundaries of GPR75 beyond any doubt. Although the conformation we observed has all the known characteristics of an active state, we cannot completely rule out the possibility that there is an even more active state or that the receptor can only adopt a single conformation, which would defy any classification of its conformation.

Along these lines, our MD simulations indicate that the conformation we observe is a stable state of GPR75, independently of the presence of the BRIL insertion. We have so far not observed a transition into a more active- or inactive-like conformation, and mutations of ECL2 promote misfolding and a non-discrete flexibility oscillating around the observed state rather than a conformational switch into a different conformation. In support of this notion, another study reporting a low-resolution structure of GPR75 observed a highly similar active-like conformation using a different fusion approach (57). Based on the accumulated structural and MD data, we assume that the conformation we observe is likely the resting state of the GPCR and the predominant state found in nature.

Despite these efforts, we cannot fully exclude the possibility that the BRIL insertion influences the conformation of our construct. At large, however, BRIL insertions into ICL3 have so far exclusively resulted in inactive or, rarely, intermediate conformations (28–32). Nonetheless, further experimental studies, such as NMR- or single-molecule FRET based approaches, might be useful to assess the conformational spectrum of unliganded wt GPR75 (reviewed in (58)).

### Constitutive activity & ECL2 self-engagement

The mutation-induced loss of ECL2 self-engagement in GPR75 leads to a loss of constitutive activity and increased conformational freedom according to our cryoEM and MD data, along with a destabilization of the receptor. A growing number of GPCRs have recently been reported to engage their own ECL2 in the orthosteric site: GPR12, GPR17, GPR21, GPR52, GPR161, and the viral GPCR BILF-1 (**Figure S4I**) (56, 59–63). These receptors delineate the different physiological effects that ECL2 self-binding can confer. As such, GPR12, GPR17, GPR21, and GPR52 possess high constitutive G-protein signaling activity that is induced by their ECL2, whereas the auto-activation of GPR161 and BILF-1 has only limited effects on signaling but is crucial for receptor expression and integrity.

In the case of GPR75, ECL2 self-engagement has a clear impact on receptor stabilization. In contrast, the amount of constitutive activity we observe for Gα_i1_ in our various assays appears low compared to e.g. GPR20 (**Figure 3B**), and we could not detect any constitutive activity for Gα_q_. Although these findings are not unprecedented as evidenced by the above examples, they do appear paradoxical considering the active-like conformation we observe as the resting state. The only other study reporting on GPR75’s constitutive activity has evaluated coupling to Gα_i1_, Gα_s_, Gα_q_, Gα_13_ and Gα_15_ as well as β-arrestin-1 and −2. Similar to our own findings, out of this limited set only Gα_i1_ and β-arrestin-2 gave a modest signal (22). In other existing studies, Gα_q_ coupling was mainly inferred from co-immunoprecipitation and by ligand-induced downstream effects in various cellular backgrounds (15, 17–20). However, it should be noted that the assays used here and elsewhere are semi-quantitative in nature and therefore not directly comparable between GPCRs.

Much like GPR75, many ECL2 self-binding GPCRs also display a low conservation among their canonical motifs and microswitches, which has been attributed to promote constitutive activity e.g. in GPR12, GPR17, and BILF1 (60, 61). An intriguing and unparalleled feature of GPR75 is the presence of a Lysine residue in the sodium binding pocket, which presumably renders the pocket unable to bind sodium ions in its (so far hypothetical) inactive state. As a result, an inactive(-like) conformation of GPR75 might be energetically unfavorable. Indicative of this, the P191A mutant appears to adopt a multitude of conformations centered around the active-like state rather than a discrete inactive state, and MD simulations over 2 µs stably retain the active-like conformation instead of diverging into a more inactive-like conformation.

### Natural ligands

We obtained no evidence of ligand-induced activity for CCL5 or 20-HETE in our set of experiments. Along those lines, several authors have recently reported negative results on both putative ligands using various assay and cell systems (3, 4, 64, 65), and the effects of CCL5 have been attributed to the presence of canonical chemokine receptors in the cellular background (20). Direct target engagement has not yet been plausibly demonstrated for either ligand, as an SPR assay monitoring the GPR75:20-HETE interaction used protein reconstituted in liposomes, which are prone to promote high affinity non-specific binding of the lipophilic 20-HETE (20). Lastly, structural comparisons of GPR75 reveal that its orthosteric pocket is quite dissimilar to those of canonical lipid- and chemokine-binding GPCRs. A lateral, allosteric 20-HETE binding site on GPR75 has been proposed, mediated by three polar residues in TM5 based on mutation studies and molecular docking (20). However, the docking has been performed *in vacuo*. It should be noted that, although we could see no direct interaction, we cannot fully exclude the possibility that either ligand might signal though yet undiscovered downstream pathways. While it is difficult to disprove binding of these ligands in absence of suitable controls, we foresee that additional experiments produced by other labs will help in clarifying this disparity.

### Conclusion, hypothesis & outlook

Based on the data accumulated in this study, namely (i) that apo GPR75 is in an active-like conformation which appears to be its resting state, (ii) that ECL2 is folded back deeply into the orthosteric site and its removal causes receptor destabilization and signaling loss, and (iii) that *bona fide* natural ligands remain elusive, we put forth the hypothesis that GPR75 might be a rare example of a constitutively active, self-activating ‘always-on’ GPCR that has no canonical inactive state and a low basal constitutive activity. Thus, GPR75 might possess no natural agonist, and instead its basal constitutive activity might be regulated through its expression levels and surface availability, or through some other unknown mechanism.

It is tempting to speculate that GPR75 might have a low inherent G-protein binding capacity or GEF activity because of its function. In line with this hypothesis, we have so far not been able to obtain a stable *in vitro* complex of GPR75 and Gα_i1._ One reason might be the unusually long ICL3 and C-terminus that crowd the cytosolic face of the receptor. These long unordered segments can modulate a GPCR’s responsiveness to its G-protein (66, 67). We tested the hypothesis that the C-terminus of GPR75 might influence its constitutive activity for Gα_i1_, but a C-terminal truncation showed no significant difference to wt GPR75 (**Figure 4C**), and we found no positive effect of C-terminal truncation on *in vitro* complex formation.

In this study we have exclusively evaluated GPR75’s function with respect to the two G-proteins Gα_i1_ and Gα_q_. Another conceivable explanation for the data we observe is that the (most) relevant physiological downstream signaling partner of GPR75 is neither of these two G-proteins, but a different effector that has not yet been identified. As such, GPR75 has one of the longest C-termini of any class A GPCR, and a study investigating naturally occurring loss-of-function mutations of GPR75 found that truncations in the C-terminus affect human BMI to equal extent as those in the rest of the molecule, seemingly contradicting our functional data with the C-terminal truncations (2). Several highly conserved stretches and phosphorylation sites within the unique, unusually long ICL3 and C-terminus might mediate interactions with GRKs, β-arrestins, scaffolding proteins, lateral interactions with other membrane proteins, combinations thereof, or some novel and yet undescribed downstream effectors. Along those lines, GPR75 has been proposed to undergo substantial constitutive internalization and recycling through an unusual dynamin-dependent, β-arrestin-independent pathway, with unknown implications for its biology (18, 64).

GPR75 has emerged as one of the most promising drug targets for obesity and related co-morbidities due to a strong genetic link in humans (2–4, 9, 16). However, its basic biology remains poorly characterized and largely controversial. Our study provides initial data shining a new light on the resting state and the activation mechanism of this unusual GPCR. However, we currently lack many of the necessary tools to further test this novel theory, and reasonable doubts remain. Future attempts to deorphanize GPR75 might provide the necessary tools to gain a better understanding of its cell biology and to unravel its downstream effectors. In addition, understanding the detailed molecular architecture and conformational space of this unusual GPCR in greater detail, e.g. by biophysical studies directly probing the conformation, will also advance our general understanding of GPCR plasticity and the role of GPCR motifs and microswitches in the understudied field of orphan GPCRs.

## Methods

The manufacturers of all important materials and instruments are listed in Table S2 (see **Supplementary Information**).

### GPR75 expression, purification, & complex formation

1L *Trichoplusia ni* High Five cells (1×10^6^ cells per mL) were infected with baculovirus encoding BRIL-fused GPR75 at a ratio of 1:555 (virus vs cell culture volume), and cells were harvested after 72h post-infection. Frozen cells were resuspended in 40 mL solubilization buffer containing 50 mM HEPES pH 7.8, 500 mM NaCl, 10 mM MgSO_4_, 10 mM L-glutamic acid (potassium salt), 10 mM L-arginine-HCl, 1% (w/v) LMNG, and 0.2% (w/v) CHS supplemented with DNase I and EDTA-free protease inhibitor. Solubilization was accomplished by rotating at 4°C for 1h, and the solubilizate was cleared by centrifugation at 65,000 g for 20-30 min. The supernatant was immobilized on 1.5 mL Streptactin XT 4Flow high capacity resin using gravity flow, washed with a gradient of solubilization to wash buffer (50 mM HEPES pH 7.8, 500 mM NaCl, 10 mM MgSO_4_, 10 mM L-glutamic acid (potassium salt), 10 mM L-arginine-HCl, 10% (w/v) sucrose, 0.01% (w/v) LMNG, and 0.002% (w/v) CHS), and eluted using a buffer containing 100 mM HEPES pH 7.8, 500 mM NaCl, 10 mM MgSO_4_, 10 mM L-glutamic acid (potassium salt), 10 mM L-arginine-HCl, 10% (w/v) sucrose, 50 mM biotin, 0.01% (w/v) LMNG, and 0.002% (w/v) CHS. The eluate was concentrated to 13.5 mg/mL and used for complex formation.

The expression and purification of anti-BRIL Fab and anti-Fab Nb are described in detail in the Supplementary Materials and Methods.

GPR75, anti-BRIL Fab, and Anti-Fab Nb were mixed in a molar ratio of 1:1.6:2 and incubated overnight at 4°C. In the case of GPR75 P191A, a molar ratio of 1:1.6:2.5 was used. The sample was then subjected to size exclusion chromatography using a Superose 6 increase column equilibrated with size exclusion buffer. The peak fractions were concentrated to 8 mg/mL and used directly for cryoEM sample preparation.

### cryoEM Sample preparation and data acquisition

2.8 uL GPR75 / anti-BRIL Fab / anti-Fab Nb complex were applied to freshly glow discharged Quantifoil R2/1 Au 300 mesh holey carbon grids. Grids were vitrified in liquid ethane using a Vitrobot Mark IV set to 4°C, 100% humidity, blot force 0, blot time 1.5s, and a wait time of 40s. Grids were first screened on a Glacios 200 kV transmission electron microscope (TEM), and data were collected on a 300 kV Krios G4 TEM equipped with a cold field emission gun, a Selectris energy filter set to a 10V slit width, and a Falcon 4i detector. Micrographs were collected at a magnification of 130,000x and a pixel spacing of 0.951 Å/px with a total dose of 40 e^-^ Å^-2^ and a nominal defocus range of −0.5 to −2 um. A total of 5,792 and 8,377 movies were collected for GPR75 wt and P191A, respectively.

### cryoEM data processing

Data processing was carried out using cryoSPARC 4.5 (**Figure S1**) (68). The micrographs were processed using patch motion correction and patch CTF estimation, followed by manual curation of the micrographs. An initial subset was used to create 2D templates, and particles were then picked using the template picker. A total of 6,665,859 particles were extracted, binned to a pixel spacing of 2.853 Å/px, and subjected to 3 rounds of 2D classification. A reference-free initial 3D model was generated using the *ab-initio* reconstruction algorithm and the particle stack was further refined as well as two rounds of heterogeneous 3D refinement. Particles were then unbinned and subjected to two additional rounds of heterogeneous 3D refinement. The purified particle stack was then subjected to a nun-uniform refinement with a mask around the whole particle including the micelle to generate a consensus map. Masks around the micelle and around BRIL + anti-BRIL-Fab + anti-Fab-Nb were generated from the lowpass-filtered consensus map in ChimeraX. These two masks were then used for focused refinements, resulting in two focused maps (GPR75/micelle and BRIL/Fab/Nb). For the masks enveloping the micelle, three consecutive local refinements were performed using a custom fulcrum located at the bottom of the orthosteric pocket. The first run was performed with a 10°/10px search radius in 0.2° increments, the second run with a 2°/2px search radius and 0.1° increment, and the final run with a 0.05° increment. A third mask enveloping all proteins was generated by molmap using a preliminary atomic model in ChimeraX, and a focused refinement was performed to generate a focused map including all proteinaceous parts of the particle. The focused and consensus maps were then locally filtered to FSC = 0.5. For the generation of a composite map, the two focused maps were aligned onto the consensus map, and the maps were combined using the volume maximum command in ChimeraX. For this purpose, the consensus map was restricted to the connecting region between GPR75 and BRIL.

### Model building and refinement

An initial model of the GPR75-BRIL chimera was generated using ColabFold (40), whereas starting models for the anti-BRIL Fab & anti-Fab Nb were based on PDB entry 7TUY. After an initial rigid body and morph fit in Coot (69), iterating rounds of manual and automated real space refinement were performed in Coot (69) and PHENIX (70), respectively.

## BRET2 assay

BRET assays were performed based on a published protocol (41–45). In short, 1.5 mio HEK293H cells in 2 mL cell culture medium (DMEM supplemented with 10% FBS, 100 U/ml penicillin and 100 µg/ml streptomycin) were seeded into 6-well plates and allowed to adhere for 3-6h. 150 ng of each G-Protein plasmid (Gα_i1_ or Gα_q_, Gβ_3_, Gγ_9_) were mixed with 150 ng of the respective GPCR plasmid. For constitutive activity, a background control containing 600 ng empty pcDNA3.1+ and a mock control containing 150 ng of each G-protein (Gα_i1_ or Gα_q_, Gβ_3_, Gγ_9_) and 150 ng empty pcDNA3.1+ instead of the GPCR were included. Cells were transfected using 2.5 uL lipofectamine 2000 per ug plasmid and incubated for 24h, washed with PBS, detached with Versene, and resuspended in cell culture medium to a density of ∼0.3-0.6 mio cells /mL. ∼30.000 to ∼60.000 cells in 100 uL were seeded into white 96-well plates in 6 technical replicates and allowed to adhere for 24h.

For constitutive activity measurements, cell culture media were replaced with 50 uL assay buffer (HBSS + 20 mM HEPES pH 7.4) and supplemented with 50 uL assay buffer containing 15 μM Prolume Purple. Plates were incubated 10 min at rt. For ligand-induced activity, the cell culture medium was replaced with 50 uL assay buffer, and 20 uL assay buffer containing 37.5 μM Prolume Purple were added. After a 5-minute incubation at rt, 30 uL ligands at 3.33x final dilution were added, plates were centrifuged briefly at 100 x g and incubated for 5 minutes at rt. The ligands were supplemented in assay buffer +0.1% BSA (final concentration 0.33%) containing a final concentration of 1% DMSO (U-46619) or 1% EtOH (20-HETE). Following incubation, plates were read in a PheraStar reader equipped with a BRET2 plus module (filters at 515-30 nm and 410-80 nm, respectively) with a gain of 3600 and using a 0.8s read interval. Seven consecutive reads were recorded and read 5 was used for data analysis. For constitutive activity, the average raw signal of the background control was subtracted from each raw data point. The BRET2 ratio was then calculated as the ratio of background-corrected GFP2 signal to the background-corrected Rluc8 signal, and the data were normalized to the average of the corresponding mock control. For ligand-induced activity measurements, no background-correction was performed, and the BRET2 ratio was calculated as before, followed by normalization against the average of the wells treated with the appropriate buffer/solvent control. Each experiment was performed in 3-5 biologically independent replicates, and values from all technical (n=3-6) and biological replicates were used for the calculation of a one-way ANOVA followed by a *post hoc* evaluation using Dunnet’s *post-hoc* test in Graphpad Prism.

## Author contributions

AML & RE designed all recombinant protein-based experiments, cryoEM, FACS, and the BRET2 assay, all of which AML performed. VG & CW designed the cAMP assay, which VG performed. JS and TK designed the Ca flux assay, which TK performed, and the quality control assays for 20-HETE together with CM. RE & AML performed the quality control assays for CCL5. AM designed and performed the MD simulations. HN provided guidance and advised the project. GB provided critical feedback. AML wrote the manuscript with help from all co-authors.

## Acknowledgements

We would like to express our gratitude to the following individuals for their contributions to this publication: Andreas Kahrs (Boehringer Ingelheim), for his helpful input and his assistance with assay development; Thomas Fox and Christoph Hoenke (Boehringer Ingelheim), for their insightful discussions; Harald Kotisch (IMP Vienna), for his operating the Titan Krios electron microscope and the cryoEM data collection; Julia Leier (Boehringer Ingelheim), for her efforts in preparing the stable CHO-K1 cell line; Bernd Guilliar (Boehringer Ingelheim) for performing the CCR5/CCL5 β-arrestin assay; Gisela Schnapp, Julian Friedl, Nikolai Prill, and Daniel Tomme (Boehringer Ingelheim), for the expression and purification of the anti-BRIL Fab & anti-Fab nanobody; Heike Rapp (Boehringer Ingelheim) for her preparation of a batch of GPR75-BRIL; Eva Martin and Tobias Claff (Boehringer Ingelheim) for guidance on setting up the BRET2 assay; Anselm Schneider and Vera Pütter (Nuvisan), for their work on initial construct screening and FSEC analysis of the GPR75 mutants; Jochen Frenzel-Pröndl (Boehringer Ingelheim), for his expertise in NMR/MS. Their input and commitment have played a crucial role in our research.

## Data availability

The cryoEM maps of the GPR75-BRIL / anti-BRIL Fab / anti-Fab Nb complex generated during the current study are available in the EMDB repository with the identifiers EMD-51796, EMD-51797, EMD-51798, EMD-51799, and EMD-51800. The coordinates of the complex are available in the PDB with the identifier 9H2F.

## Additional files

Supplementary Information

