## Supplementary Information for "The CryoEM Structure of Human GPR75: Insights into ECL2-Mediated Activation"

This PDF file includes:

Supplementary Materials and Methods

Figures S1 to S7

Tables S1 to S2

SI References

### Supplementary Materials and Methods

#### Constructs

The codon-optimized human GPR75 fused with an N-terminal signal peptide followed by a FLAG tag, a BRIL insertion between residues A233$^{5.69}$ and L311$^{6.25}$, a C-terminal truncation after residue 398 and a PreScission-cleavable GFP-TwinStrep-10xHis-tag was cloned into pFastBac1. The anti-BRIL Fab light and heavy chain (1) were synthesized by GeneArt (Thermo Fisher Scientific, Darmstadt, Germany), and cloned separately into a pTT5 vector controlled by a CMV promoter. Anti-Fab Nb with a secretion signal and a TEV-cleavable N-terminal His tag (2) was synthesized by GeneArt and cloned into a pET24 vector. pcDNA3.1+ plasmids containing the constructs for BRET2 assays were synthesized by GeneArt. Except for the ECL2 mutants, all GPCRs were wt sequences, and the sequences of $G\alpha_{i1}$-RLuc8, $G\alpha_q$-RLuc8, $G\beta_3$, and $G\gamma_9$-GFP2 were based on a published protocol (3).

#### Generation of a stable CHO-K1 cell line expressing GPR75

CHO-K1 cells were transfected with NruI-cut, linearized pcDNA3.1-GPR75 plasmid by use of FuGENE HD transfection reagent in a 6-well culture plate following manufacturers' instructions. Two days after transfection cells were split into selective medium Ham´s F12 + 10% FBS + 400 µg/ml geneticin to establish a stable pool of transfected cells. From day six post transfection, clonal cell lines were established by limited dilution seeding and continuous maintenance in selective medium. Expression of GPR75 was verified on the RNA level in a qRT-PCR and on protein level by FACS analysis.

**Protein expression, purification, & complex formation**

GPR75 mutants

For nanoDSF experiments, GPR75 ECL2 mutants were purified analogously to GPR75-P191A. These samples were snap-frozen and stored at -80°C until further use. For cryoEM GPR75 P191A was expressed and purified analogous to GPR75 wt, except that immobilization to the affinity resin was accomplished by batch binding, rotating 1 h at 4°C, 15 rpm.

Anti-BRIL Fab

125 µg of each heavy and light chain plasmid in 32 mL OptiPRO SFM were mixed with 188 µL cold TransIT Pro and added to 250 mL CHO-3E7 cells in BalanCD Transfectory CHO medium + 4mM GlutaMAX at a density of 2 x$10^6$ cells/mL. Fab fragments were expressed for 7-10 days at 37°C, 5% $CO_2$, 120 rpm and fed with 25 mL Irvine Feed Transfectory Supplement and 625 µL Anti-Clumping reagent after 24 h. After 6 days, 50 mL CHO CD Efficient Feed B were added. Cell supernatants were harvested by centrifugation at 4,500 rpm for 20 min. The cell culture supernatant was supplemented with 30g of Sartoclear Dynamics Lab Filter Aid to supernatant and sterile filtered. 2 mL KappaSelect beads per Liter cell culture (1000rpm, 5min) were added and incubated overnight at 4°C on a roller mixer. The beads were transferred into a flow cartridge, washed with PBS until A280 dropped below 0.01 mAU, and eluted using a buffer containing 30mM sodium acetate pH 3,0 and 100mM NaCl. The eluate was neutralized by adding 10% (v/v) of a buffer containing 300mM sodium acetate pH 9,0 and 100mM NaCl. The sample was then dialyzed 3x overnight at 4°C in 4 L PBS pH 7.4, concentrated to 1.5 mg/mL, snap-frozen and stored at -80°C. Before complexation with GPR75, the anti-BRIL Fab was buffer-exchanged into size exclusion buffer (20 mM HEPES pH 7.8, 150 mM NaCl, 10 mM $MgSO_4$, 10 mM glutamic acid (potassium

salt), 10 mM L-arginine-HCl, 0.004% (w/v) LMNG, and 0.0008% (w/v) CHS) using a PD-10 column and concentrated to 7.4 mg/mL.

Anti-Fab Nb

The anti-Fab nanobody was expressed periplasmatically in *E.coli* BL21 (DE3) for 24 h at 20°C in LB medium. Cells from 1L were harvested and resuspended in 200 mL of a buffer containing 20 mM Tris-HCl pH 7,5 and 150 mM NaCl supplemented with DNAse I and Complete EDTA free protease inhibitor. Cells were lysed using a microfluidizer at 700 bar in 10 cycles. Cell debris were removed by centrifugation at 45,000 x g, 35 min, and the supernatant was incubated with 6 mL Ni-NTA resin for 1.5 h at 4°C while rotating. Beads were then washed with 200 mL of a buffer containing 20 mM Tris-pH 7,5, 500 mM NaCl, and 30 mM imidazole, and protein was eluted using 5 CV of a buffer containing 20 mM Tris-pH 7,5, 500 mM NaCl, and 250 mM imidazole. The sample was supplemented with 1 mM DTT and 10% (v/v) glycerol, and the His-tag was cleaved over night by adding 1.5 mg TEV protease while dialyzing against 4 L of a buffer containing 20 mM Tris-HCl pH 7,5, 200 mM NaCl, and 2,5 % (v/v) glycerol. The sample was concentrated and applied to a Superdex 75 column equilibrated with a buffer containing 20 mM Tris-pH 7,5 and 150 mM NaCl. Peak fractions were pooled, concentrated, and stored at -80°C. Before complex formation, the anti-Fab Nb was buffer-exchanged into size exclusion buffer using a NAP-5 column and concentrated to 8.6 mg/mL.

**STAGS analysis of GPR75 conformation**

STAGS (4) was used with standard parameters and the default training set, using as input either our complete experimental structure including BRIL, the structure with all

BRIL residues removed, and a version of the structure in which the wt ICL3 from the prediction available at the AlphaFold database was grafted manually onto our experimental structure. These three methods returned probabilities of 98%, 98%, and 97% for an active-like conformation, respectively.

**Structural comparison between GPR75 and β1-AR**

Alignment between GPR75 and the active (PDB 7BU7) and inactive (7BVQ) conformations of β1-AR were performed using USalign with standard settings (5). For visualization, Alignment was performed using matchmaker in ChimeraX (6).

**MD simulations**

The experimental structure of GPR75-BRIL was used as starting point for all MD structures. Point mutations and/or ECL2 ejections were introduced manually in Coot (7). The wt ICL3 was grafted onto the experimental structure by alignment with the GPR75 AlphaFold2 model (AF-O95800-F1-v4) in Coot. The structures were embedded in a lipid bilayer consisting of POPC with 15% cholesterol using the packmol-memgen program in Ambertools 21 (8). Amber ff99SB-ILDN force field was used for protein parameters in GROMACS 2021 (9). SLIPIDS parameters were used for lipids (10)**.** The simulation systems were solvated with the TIP3P (11) water model. Counter ions were added to neutralize the net charge of simulation systems. Hydrogen mass repartitioning scheme (HMR) (12, 13) was used to achieve a 4 fs integration timestep for all simulations; hydrogen masses, except those of waters, were increased to 3 amu. Hydrogen motions were constrained using the LINCS algorithm (14, 15). In all cases, the simulation temperature was set to 298.15 K. A simulation pressure of 1 atmosphere was maintained using Berendsen barostat (16) during equilibration with a time constant of 1 ps, followed by the Parrinello-Rahman barostat (17) with a time constant of 2.0 ps for production simulations. A cut-off of 1 nm was used for short range interactions, and long-range electrostatics were handled via PME (18, 19). The simulations system was minimized. Next, five step equilibration was used in which protein backbone atoms, non-hydrogen ligand atoms and lipid head groups were restrained using the following force constants: 5, 2.5, 1, 0.5, and 0.1 kcal per mol per Å$^2$. Then, an additional equilibration step was performed with position restrains only applied to the protein backbone and non-hydrogen ligand atoms of 0.1 kcal per mol per Å$^2$. All simulations were conducted for 2 µs with three replicates, first 100 ns were treated as equilibration phase, based on the area per lipid property and all analysis was performed on later 1900 ns. The trajectory analysis was performed using the GROMACS tools (9) and the MDAnalysis package (20).

**nanoDSF**

For the measurements of ECL2 point mutants, all samples were diluted to 0.4 mg/mL in a buffer containing 100 mM HEPES pH 7.8, 500 mM NaCl, 10 mM $MgSO_4$, 10 mM L-glutamic acid (potassium salt), 10 mM L-arginine-HCl, 10% (w/v) sucrose, 50 mM biotin, 0.01% (w/v) LMNG, and 0.002% (w/v) CHS.

For measurements of ligand-induced thermostabilization, purified GPR75 and GPR52 were buffer-exchanged into a buffer containing 20 mM HEPES pH 7.8, 0,005% (w/v) LMNG, 0,001% (w/v) CHS, and 150 mM NaCl. Lyophilized CCL5 was reconstituted directly in this buffer, as well. 20-HETE (stored in ethanol) was dried quickly and carefully with nitrogen gas and reconstituted in the same buffer. The two GPCRs (final concentration 3 μM) were mixed with either CCL5, 20-HETE (6 and 9 μM final concentration. respectively) or buffer and incubated 5 min at rt in the dark prior to the measurement.

nanoDSF measurements were performed using a Prometheus NT.48, melting at 1°C/min from 20 to 95°C in high sensitivity capillaries, recording fluorescence signals at 330 and 350 nm. The runs were performed at 75% (ECL2 mutants) or 90% (ligand experiments) excitation power. For each run, the first derivative of the 350 nm signal or the 330/350 nm ratio was calculated. Traces from 3-6 technical replicates were averaged and graphed in GraphPad Prism.

**FSEC analysis of protein expression**

FSEC analysis was performed at Nuvisan GmbH (Berlin, Germany). Point mutants were expressed in 2 mL ESF cells for 72 h at an MOI of 1 and solubilized in a buffer containing 50 mM Tris pH 7.5, 150 mM NaCl, 1% LMNG, and 0.1% CHS. Samples were cleared by ultracentrifugation, filtered, and applied on a Dionex HPLC equipped with a Superose 6i column. Fluorescence counts were recorded and plotted using GraphPad Prism.

**cAMP assay**

CHO-K1 clones stably overexpressing GPR75 were brought into suspension with non-enzymatic cell dissociation solution and neutralized with growth medium containing Ham's F-12 (Kaighn's) supplemented with 10% HI-FBS and 1% Pen-Strep. Cells were seeded either at 750/1500/3000 cells per well in a 384 well plate in 25µL growth medium. The next day the cells were treated with Forskolin at various concentrations for 1 hour in a humidified 37C, 5% CO2 incubator. cAMP was detected according to the kit protocol. For each cell density, three technical replicates were recorded.

**Calcium flux assay**

The Calcium flux assay was performed using the FLIPR Calcium 6 assay kit with CHO-K1 cells transiently overexpressing GPR75. For transient transfection, 2.0 $x10^5$ CHO-K1 cells per well were seeded on a 6-well culture plate in Hams´F12 medium containing 10% HI FBS and 1% Pen/Strep and incubated overnight at 37°C, 5% $CO_2$, humidified atmosphere. 2 µg wt GPR75 in pcDNA3.1 plasmid were diluted in 150 µL OptiMEM, mixed with 6 µL FuGENE HD reagent and incubated 15 min at RT before transfection. 24 h after transfection, cells were suspended in Ham’s F12 medium supplemented with 10% FBS, and 1.5 $x10^4$ cells were seeded into each well of a black clear-bottom 384 well plate. After 24 h, cells were starved for 2 h by exchanging the medium to 20 µL serum-free Ham’s F12. Following the starvation period, the medium was exchanged to 20 µL assay buffer (HBSS + 5 mM HEPES, pH 7.4 and 0.1% (w/v) BSA). 20 µL of assay buffer supplemented with Calcium-6 dye according to the manufacturer’s recommendations and 5 mM Probenecid were added to each well and incubated for another 2 h at 37°C. The assay was run on a FLIPR Tetra (Molecular Devices) equipped with a CCD camera. Following a 10s baseline measurement, 10 µL of the appropriate 5x compound dilution in assay buffer + 0.5% EtOH were added to the plate and the changes in fluorescence were read in real-time over a period of 118 s. For each data point, the difference between the minimal and maximal fluorescence was plotted against the corresponding ligand compound concentration. Data were collected in three technical replicates. A non-linear fit was plotted for each sample.

**PathHunter CCR5 assay**

The PathHunter eXpress β-arrestin assay was performed according to the manufacturer's recommendations. Briefly, cells were thawed in 12 mL pre-warmed AssayComplete cell plating reagent, and 100 µL per well of the resulting cell suspension were seeded into a 96-well plate. Plates were incubated for 48 h. 10 µL of either CCL5 or CCL3 were added at 11x the desired final concentration, and the plates were incubated for another 90 min. 55 µL of working detection solution were added to the wells, and the plates were incubated for 1 h at rt in the dark. Assay plates were read on an EnVision Multilabel Plate Reader at 0.1 s/well. Data were collected in two technical replicates, plotted, and fitted in GraphPad Prism.

**20-HETE quality control**

20-HETE (Cayman) was stored in EtOH, -20°C, in the dark under argon or nitrogen. Final dilutions were prepared freshly before each experiment and used as rapidly as possible.

*Rat aortic ring constriction assay*

The assay was performed by PhysioStim (Castres, France). For the preparation of aortic rings, the heart of male rats with the aorta was rapidly excised and placed into fresh saline solution previously equilibrated for 30 minutes with a 95% $O_2$ / 5% $CO_2$ gas mixture. The rat thoracic aorta was quickly dissected, cleared of connective tissue, and cut into cylindrical segments. Two stainless steel triangles were passed through the lumen of aortic rings, one of which was fixed to the wall of a temperature-controlled emkaBATH4 organ bath whereas the other one was attached to a force transducer. The prepared aortic rings were mounted into the organ bath containing 20 mL of a solution (118.0 mM NaCl, 25.0 mM $NaHCO_3$, 1.2 mM $KH_2PO_4$, 1.2 mM $MgSO_4$, 2.5 mM $CaCl_2$, 4.7 mM KCl, and 11.1 mM Glucose) which was continuously gassed with 95% $O_2$ / 5% $CO_2$ and continuously maintained at +37.0 ± 0.5°C. The aortic rings were stretched to optimal length (approximately 1 x g). The organ bath solution was replaced each 15 minutes during a stabilization period of at least 30 minutes before two successive challenges with 60 mM KCl. Vessels were then allowed to stabilize at least 30 minutes and endothelial integrity was assessed by using 1 μmol/L acetylcholine to induce vessel relaxation after a pre-constriction with 1 μmol/L phenylephrine. The organ bath solution was then replaced each 15 minutes during a stabilization period of at least 30 minutes. During the measurement period, 20-HETE was added in cumulative steps to the appropriate final concentrations, starting at 3 μM. At each experimental step, the isometric tension was continuously measured for at least 5 min and until a stable effect was obtained, up to a maximum of 30 min, to detect any vasocontractile effect. The isometric tension was recorded and analyzed using IOX. Data were collected in two independent experiments.

### Supplementary Figures

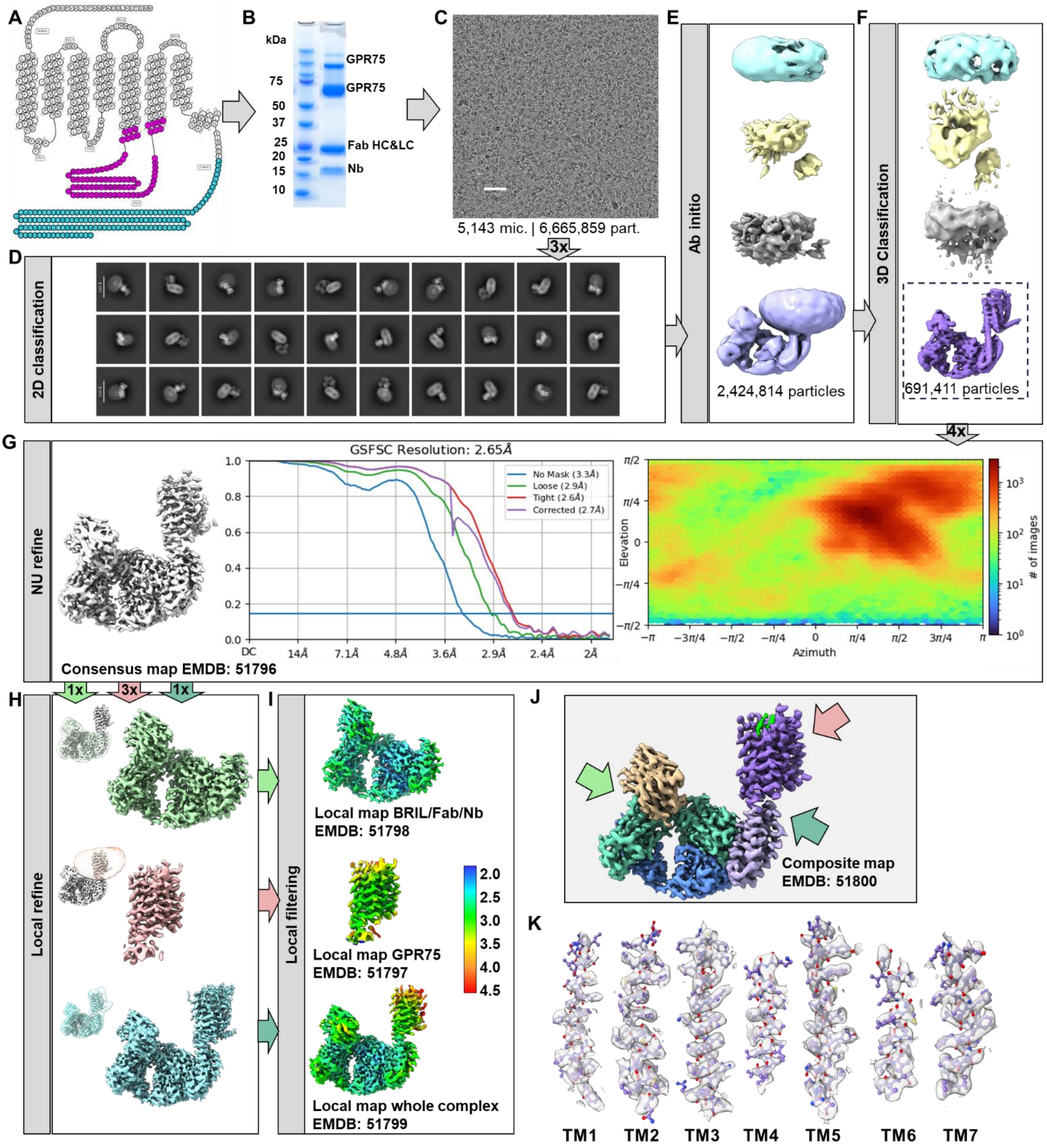

**Figure S1 | CryoEM workflow. Related to Figure 1. A** Snake plot showing the GPR75 sequence (21). ICL3 residues (magenta) were replaced by BRIL, C-terminal residues (teal) were removed. **B** SDS-PAGE of the cryoEM sample. **C** Representative micrograph recorded at -2.3 Å measured defocus. **D** Representative 2D classes after 3 iterations. **E** Reference maps generated by *ab-initio* reconstruction. These maps were used as references for subsequent 3D classification. **F** Maps after 4 rounds of 3D classification. **G** Non-uniform refinement map, including FSC curve and angular distribution plot as reported by cryoSPARC (22). **H** Local refinement maps using the masks indicated in the small inlays. For GPR75, 3 consecutive local refinements with increasingly fine sampling were performed. **I** Local resolution and filtering of the focused maps. All maps are colored according to the key on the left side of the panel. **J** Composite map generated in ChimeraX (6) from the three local maps, contoured at a level of 0.29. **K** Density around the individual helices, contoured at a level of 0.26.

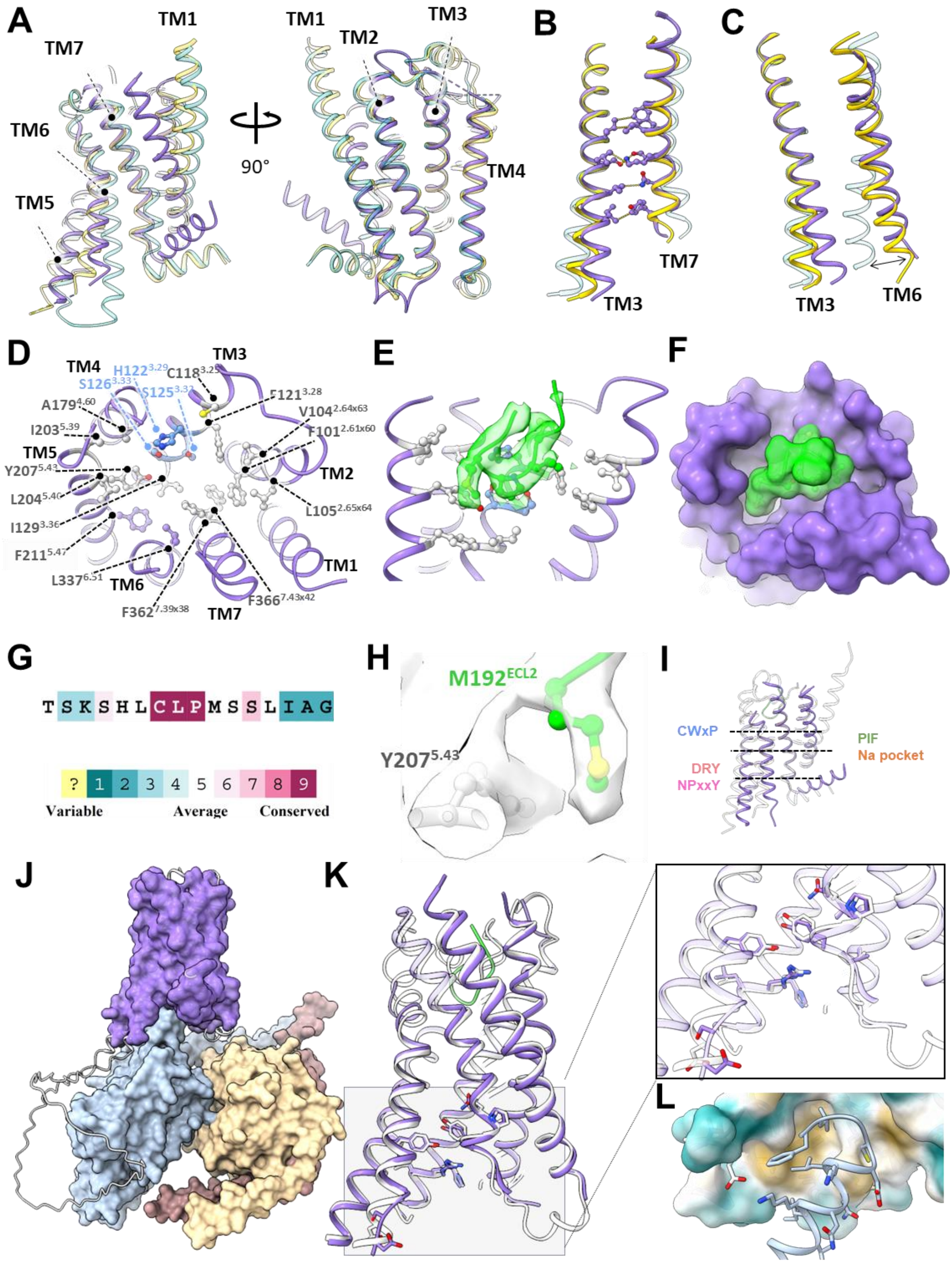

**Figure S2 | Conformational state and orthosteric pocket of GPR75. Related to Figure 1. A** Comparison of the GPR75 tertiary structure to the active (PDB 7BU7, yellow) and inactive (7BVQ, teal) states of β1-AR. **B** Interaction network between TM3 and TM7. **C** Conformation of TM6 relative to TM3. **D** Top view of the orthosteric pocket showing the locations of all individual residues. GPR75 is colored in purple, hydrophobic residues lining the orthosteric pocket are colored in gray. Polar residues are colored in blue. ECL2 is omitted for clarity. **E** Side view of the orthosteric pocket showing the cryoEM density of ECL2 (green surface) rendered at a level of 0.24. TM6&7 are omitted for clarity. **F** Surface representations of GPR75 shown from the top, with ECL2 colored lime. The bulk of ECL2 fills up the orthosteric site of GPR75. **G** Sequence conservation of the 'CLPM' motif, generated with ConSurf (23). **H** Potential alternative conformation of $M192^{ECL2}$. **I** Position of the canonical motives displayed in Figure 3A-F. **J-L** The active state of GPR75 (white) was modeled using ColabFold (24) in complex with human $G\alpha_{i1}\beta_1\gamma_2$ (blue, yellow, orange), and the complex was superposed with the experimental coordinates of GPR75 (purple) in ChimeraX. **J** Overview of the superposition, showing the experimental structure of GPR75 (purple) and the computed structures of $G\alpha_{i1}\beta_1\gamma_2$. The computed GPR75 model (including ICL3) is shown as white ribbons. **K** The tertiary structure and residues of the G-protein binding pocket of GPR75 were superposed with the computed active state. Zoom-in shows a detailed view of the superposition in the G-protein binding cavity. **L** The superposed, calculated C-terminus of $G\alpha_{i1}$ fits into the experimentally determined pocket without major clashes. The experimental structure is shown as surface and colored by hydrophobicity (gold=hydrophobic, teal=hydrophilic). The calculated C-terminus of $G\alpha_{i1}$ is shown in light blue.

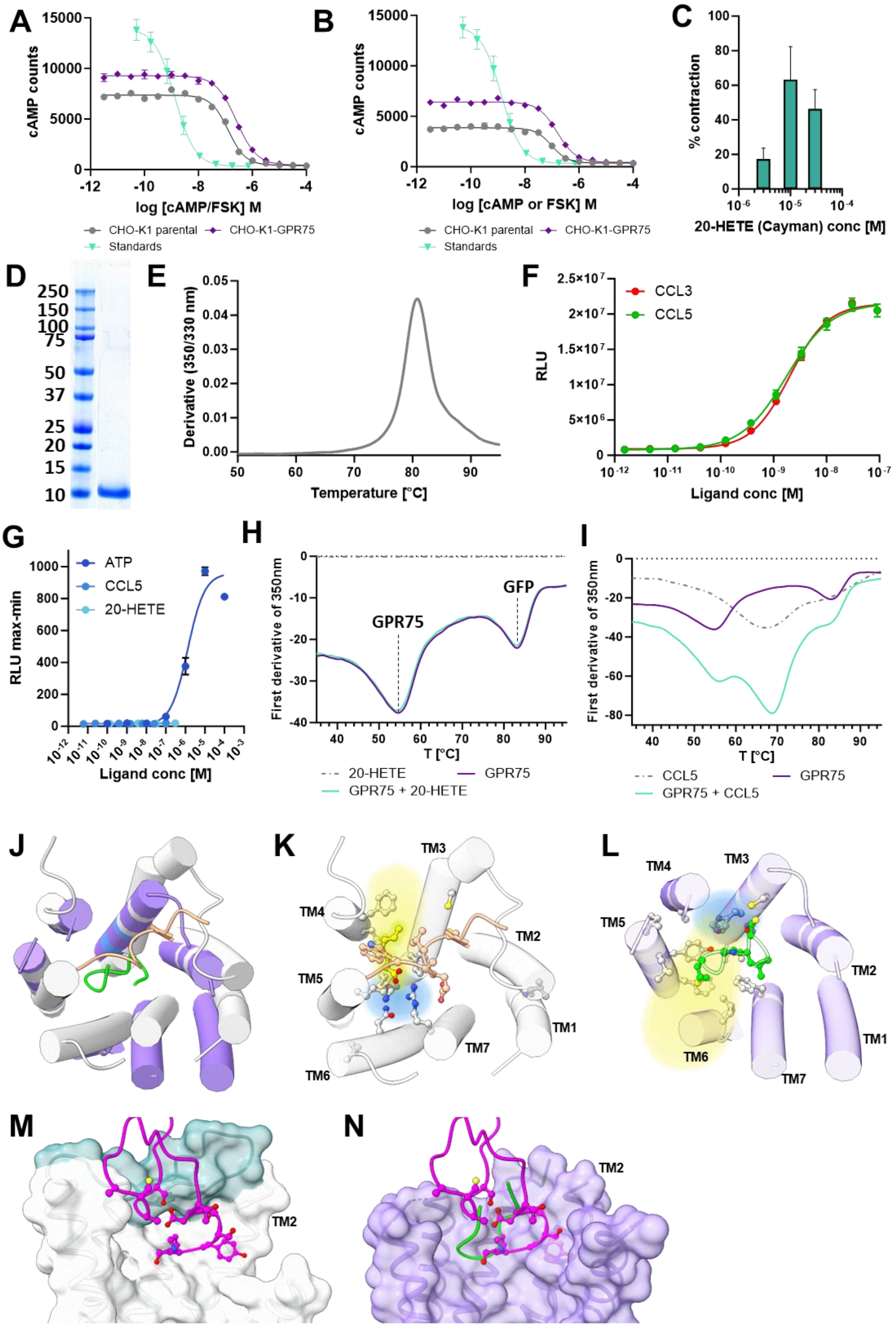

**Figure S3 | Quality control of GPR75 ligands and ligand-induced activation of GPR75. Related to Figure 3. A,B** Repetition of cAMP assay (Fig. 3C). Assay conditions were identical, except that 1,500 (A) and 3,000 (B) cells per well were used. **C** Rat aortic ring constriction assay with increasing concentrations of 20-HETE. **D** SDS-PAGE showing the purity of CCL5. **E** nanoDSF of CCL5 (in PBS) shows the protein is folded. **F** β-arrestin assay showing the efficacy of CCL5 and CCL3 (control chemokine) on the PathHunter CCR5 cell line (DiscoverX). **G** $Ca^{2+}$ flux assay with CHO-K1 cells transiently expressing GPR75. Data are mean ± SEM of three technical replicates. **H-I** nanoDSF of purified recombinant GPR75 (3 μM) in the presence and absence of 20-HETE (9 μM, H) or CCL5 (6 μM, I). **J-L** Comparison of the GPR75 orthosteric pocket (L) to the pocket of ligand-bound FFAR1 (PDB 8EIT, J-K). **M** CCL5 (magenta) shown in complex with CCR5 (PDB 7F1R). The CCR5 ECL2 is colored teal. **N** CCL5 from the CCR5 complex structure (magenta, 7F1R) clashes with GPR75 (purple) TM2 & ECL2 (lime) when the GPCRs are superposed. Positions of TM2 are indicated. (M-N) TM 5&6 are omitted for clarity.

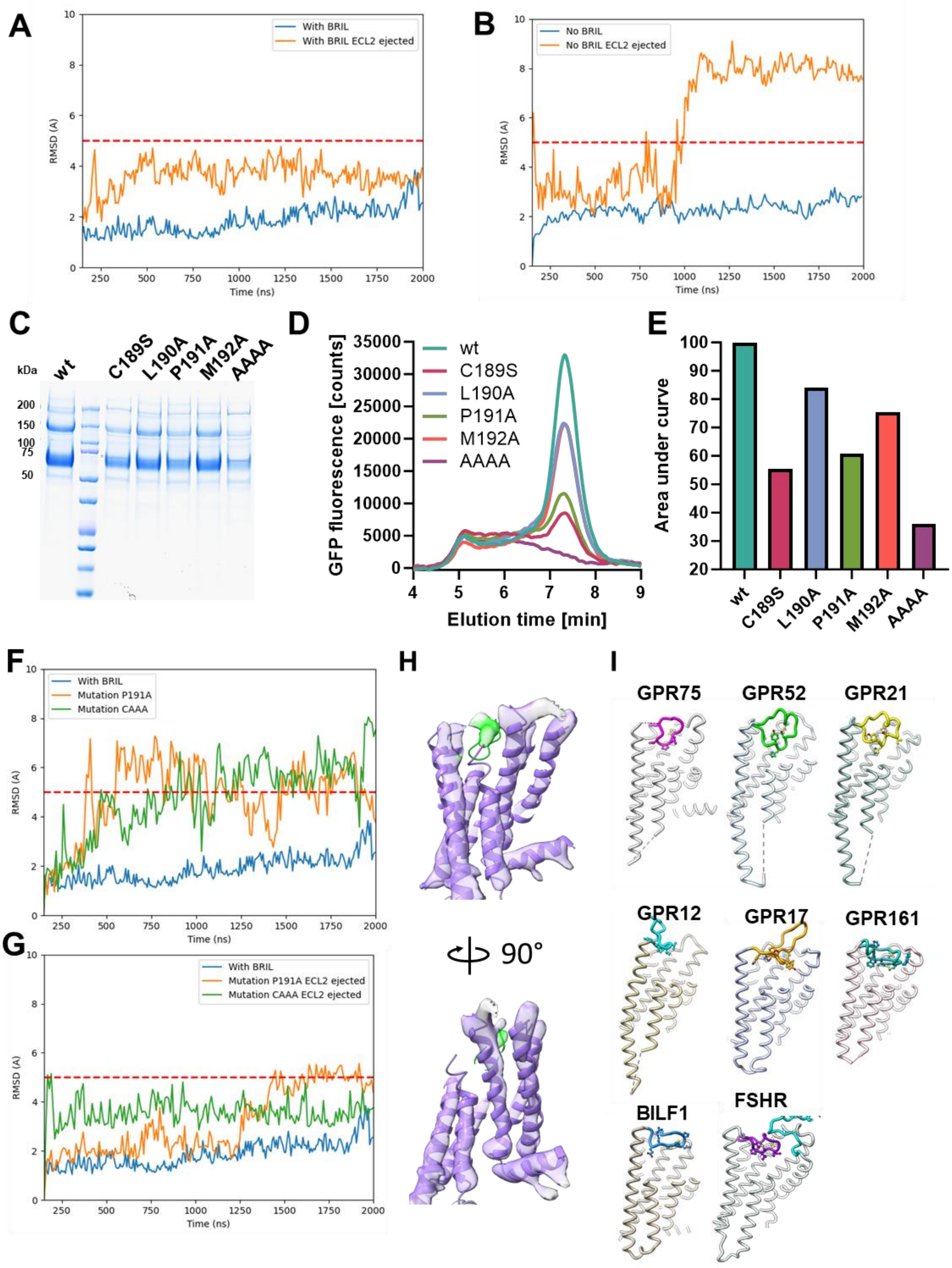

**Figure S4 | Effects and comparison of the GPR75 ECL2. Related to Figure 4. A,B** 2 µs MD simulation of GPR75 with and without artificially ejected ECL2 in a lipid bilayer, either including the BRIL insertion (A) or wt ICL3 (B). Blue: reference rmsd trace of a simulation using the experimentally observed structure. **C** SDS-PAGE showing the purity of the recombinant ECL2 mutants. **D-E** FSEC performed with ECL2 mutants of the GPR75-BRIL construct after overexpression in Hi5 insect cells. B: FSEC chromatogram. C: Area under curve, indicating the relative yield after expression and solubilization. **F-G** 2 µs MD simulation of GPR75 ECL2 mutants in a lipid bilayer, either by point mutation of the observed structure (F) or by point mutation in combination with artificial ECL2 ejection (G). Blue: reference rmsd trace of a simulation using the observed structure. **H** The GPR75 wt coordinates (ribbons) docked into the P191A map. **I** Comparison of ECL2 self-binding GPCRs. ECL2s are colored. PDB codes: 9H2F (GPR75), 6LI3 (GPR52), 8HMV (GPR21), 7Y89 (GPR17), 7Y3G (GPR12), 7JHJ (BILF-1), 8KH4 (GPR161), 8I2G (FHSR).

| | | | |
|---|---|---|---|
| 306 | 1 | GPR75_human_O95800 | ATLATLKT------SKSHLCLPMSSLIAGKGK-AILSLYV |
|  | 2 | UniRef90_F6XQB5_start_1_end_540_Evalue_0 | ATLATLKT------SKSHLCLPMSSLIAGEGK-AILSLYV |
|  | 3 | UniRef90_A0A287AG85_start_1_end_540_Evalue_0 | ATLATLKT------SKSHLCLPMSSLIAREGK-AILSLYV |
|  | 4 | UniRef90_A0A836AJ96_start_1824_end_2363_Evalue_0 | ATLATLKT------SKSHLCLPMSSLIAGEGK-AILSLYV |
|  | 5 | UniRef90_M3WIQ7_start_1_end_514_Evalue_0 | ATLGTLKT------SKSHLCLPMSSLMAGEGR-AILSLYV |
|  | 6 | UniRef90_D2GY88_start_1_end_540_Evalue_0 | ATLGTLKT------SKSHLCLPMSSLTAGEGK-AIVSLYV |
| 307 | 7 | UniRef90_Q6X632_start_1_end_540_Evalue_0 | ATLATLRT------NKSHLCLPMSSLMDGEGK-AILSLYV |
|  | 8 | UniRef90_UPI0010A1E234_start_4_end_533_Evalue_0 | ATLATLRT------SKSHLCLPMSSLMAGEGK-AILSLYV |
|  | 9 | UniRef90_UPI00187A681B_start_1_end_540_Evalue_0 | ATLATLKT------SQSHLCLPMSSLIAGDGK-AVLSLYV |
|  | 10 | UniRef90_G1PZ15_start_1_end_536_Evalue_0 | ATVATLKT------RKSHLCLPMSSLMAGEGK-AVLSLYV |
|  | 11 | UniRef90_UPI000332EF8E_start_9_end_546_Evalue_0 | ATLATLRT------NKSHLCLPMSSLMAGEGK-AVLSLYV |
|  | 12 | UniRef90_UPI000333BA05_start_1_end_540_Evalue_0 | ATLATLKT------SKSHLCLPMCSVIAGEGR-AVLSLYV |
|  | 13 | UniRef90_UPI0011B018A3_start_4_end_535_Evalue_0 | ATLGTLKT------SKSHLCLPMSSLMAGEGR-AILSLYV |
|  | 14 | UniRef90_A0A286X9Q4_start_1_end_530_Evalue_0 | ATLATLKT------SKSHLCLPMSSLMAREGR-AILSLYV |
|  | 15 | UniRef90_A0A7J7VUR1_start_1_end_537_Evalue_0 | ATVATLKT------RTPPLCLPMSSLAAGEGR-AVLSLYV |
|  | 16 | UniRef90_UPI0003F08F6A_start_1_end_541_Evalue_0 | ATLATLKT------SKSHLCLPMSSLIAGEGK-AVLSLYV |
|  | 17 | UniRef90_UPI0007A72E4E_start_1_end_539_Evalue_0 | ATLATLKT------SKSHLCLPMSSVIAGGGR-AVVSLYV |
|  | 18 | UniRef90_F7ACL6_start_1_end_537_Evalue_0 | AALATLKT------SKSRLCLPMSSLITGEGK-TILSLYV |
|  | 19 | UniRef90_A0A7L2JU52_start_14_end_508_Evalue_1.1e-213 | ATLVTLRT------RGSRFCLPMASLSSGEGK-LILYLYV |
|  | 20 | UniRef90_A0A7J5ZLZ3_start_16_end_547_Evalue_1.1e-217 | AALVTLRAYP----RSAGPCLPHFGLTGNRAK-VVLYVYL |
|  | 21 | UniRef90_UPI001C64FE5D_start_196_end_708_Evalue_1.1e-260 | ATMATLRT------RGSRLCLPMSSFARGDGK-VILYLYV |
|  | 22 | UniRef90_A0A8C2W5W6_start_1_end_534_Evalue_1.1e-285 | ATLGTLKT------GKSHLCLPMSSLVAREGR-AILSLYM |
|  | 23 | UniRef90_UPI001403062B_start_22_end_516_Evalue_1.2e-198 | ATLSAMTKY-----HSSKICVPLLGLINEDGK-AILYTYV |
|  | 24 | UniRef90_A0A672TRA6_start_1_end_429_Evalue_1.2e-211 | ATLATLKT------RRSRLCLPMSSFASREGK-VILYLYV |
|  | 25 | UniRef90_A0A087YQ19_start_13_end_547_Evalue_1.2e-215 | AALLTMRAYP----RRDGPCLPHFGLGSGQAR-VVLYVYL |
|  | 26 | UniRef90_A0A7K7ICF1_start_14_end_513_Evalue_1.2e-220 | ATLATLRT------RGSRLCLPMASPPRGQGK-LILYLYV |
|  | 27 | UniRef90_A0A8C5BSR9_start_33_end_553_Evalue_1.2e-220 | AALLTMRAYP----RKDGPCLPHFGLSGSRAR-VVLYAYM |
|  | 28 | UniRef90_A0A7K8VS48_start_3_end_492_Evalue_1.2e-255 | ATLATLRA------HSSRLCLPMASFASGQGK-IILYLYV |
|  | 29 | UniRef90_A0A6P8Q3P0_start_27_end_537_Evalue_1.2e-270 | ASLATLRT------RKSSICLPVYSLASGEGK-AVLYLYV |
|  | 30 | UniRef90_F1ND05_start_24_end_536_Evalue_1.2e-273 | ATMATLRT------HGSRLCLPMSSFARGDGK-VILYLYV |
|  | 31 | UniRef90_A0A7L2QLE1_start_14_end_477_Evalue_1.3e-204 | ATLAALRT------RGSRLCLPVASLPRARGK-LILYLYL |
|  | 32 | UniRef90_A0A852J8K3_start_15_end_544_Evalue_1.3e-266 | ATLATLKT------RGSRLCLPMSSFASDEGK-IILYLYV |
|  | 33 | UniRef90_A0A7K8SXI7_start_34_end_543_Evalue_1.3e-273 | ATLATLKT------RGSRLCLPMSSFTSGEGK-IILYLYV |
|  | 34 | UniRef90_A0A2I0M9A2_start_1_end_332_Evalue_1.4e-159 | ----------------------MSSFAGGEGK-IILYLYV |
|  | 35 | UniRef90_A0A7K4WLP5_start_14_end_503_Evalue_1.4e-211 | ATLATLRA------RGSRRCLPMASLAIGQGR-LILYLYI |
|  | 36 | UniRef90_A0A4W4F2I0_start_31_end_551_Evalue_1.4e-216 | AALVTLRVYP----SSTGPCLPHFGLAGRPAQ-VVLCVYL |
|  | 37 | UniRef90_A0A8C9W592_start_33_end_549_Evalue_1.4e-223 | ATLLTLHAHP----WGSRPCLPRFGLAGGDGK-VTLCLYL |
|  | 38 | UniRef90_W5NLF8_start_31_end_533_Evalue_1.4e-264 | ATLVTLRTYP----WKSRVCLPHFGLIGGDGK-IILYLYL |
|  | 39 | UniRef90_UPI00053043E7_start_2_end_368_Evalue_1.5e-185 | ------KP------PAPRLCLPMSSFASGEGK-IILYLYV |
|  | 40 | UniRef90_A0A401SVN0_start_39_end_533_Evalue_1.5e-201 | ATLSAMTKY-----HSSKICVPLLGLINEDGK-AILYTYV |
|  | 41 | UniRef90_A0A7K7T670_start_13_end_484_Evalue_1.5e-207 | ATLATLRT------RGSRRCLPMASLAGGQGR-LVLYLYL |
|  | 42 | UniRef90_A0A401SSV5_start_10_end_538_Evalue_1.5e-218 | ATLATMRTYP----ERSKVCVPLFGCIKEDDK-IILYLYV |
|  | 43 | UniRef90_A0A8C4JJJ0_start_1_end_444_Evalue_1.5e-242 | ATLATLKT------RRSRLCLPMSSFASGEGR-VILYFYV |
|  | 44 | UniRef90_A0A7K6RYJ2_start_3_end_502_Evalue_1.5e-261 | ATLATLKT------RGSRLCLPMSSFAGGEGK-LILYLYV |
|  | 45 | UniRef90_A0A7K5HGS4_start_1_end_502_Evalue_1.5e-264 | ATLATLRS------RSSRLCLPMSSFAGSQGK-VILYLYV |
|  | 46 | UniRef90_A0A7L2HUQ2_start_3_end_512_Evalue_1.5e-275 | ATLATLKT------HGSRLCLPMSSFASGKGK-IILYLYV |
|  | 47 | UniRef90_A0A6P7XJL5_start_26_end_538_Evalue_1.5e-286 | ATLATLRT------RKSRSCLPMVSLLSGEGK-VVLYLYV |
|  | 48 | UniRef90_UPI0014025B9A_start_25_end_472_Evalue_1.6e-151 | AILTTFRTYP----DSSRVCVPLFGCSNRDGR-VILYLYV |
|  | 49 | UniRef90_UPI001F07A78A_start_15_end_520_Evalue_1.6e-198 | ATLLTVRAYA----RRDGPCLPHLGPAGARAR-VVLYVYL |
|  | 50 | UniRef90_UPI00074FF0FF_start_92_end_627_Evalue_1.6e-294 | ATLATMRT------HKSRLCLPISSLVSGEGK-IIPYLYV |
|  | 51 | UniRef90_UPI0015B159A8_start_30_end_554_Evalue_1.7e-226 | AALFTLRTYP----WRSRPCLPHAGLVAVEGK-AVLYLYL |
|  | 52 | UniRef90_UPI0010A8E366_start_3_end_407_Evalue_1.8e-123 | AALATLRP------ARSRLCLPLAGPPGGGGT-TVLCLYA |
|  | 53 | UniRef90_A0A7K9KCS9_start_14_end_507_Evalue_1.8e-214 | ATLATLRT------RGSRLCLPMASLPSGQGK-LILYLYV |
|  | 54 | UniRef90_UPI001C091AEF_start_33_end_568_Evalue_1.8e-220 | AMLVTLRAYP----RSAGPCLPHFGLTGNRAK-VVLYVYL |
|  | 55 | UniRef90_Q6GM03_start_7_end_511_Evalue_1.8e-284 | ATLATLRT------RKSRICLPMYSLINGEGK-IILYLYV |
|  | 56 | UniRef90_UPI00053CE52B_start_128_end_528_Evalue_1.9e-210 | ATLATLKT------HGSRLCLPMSSFASGEGK-IILYLYV |
|  | 57 | UniRef90_A0A670HT06_start_22_end_548_Evalue_1.9e-287 | ATLATMRT------QKSRLCLPISSLVIGEGK-VVPYLYV |
|  | 58 | UniRef90_A0A7K4R454_start_9_end_501_Evalue_1e-205 | ATLAALRA------RGSRCCLPVASLAGGRGG-LALYLYL |
|  | 59 | UniRef90_A0A093H6W6_start_1_end_393_Evalue_1e-220 | ATLATLKT------RRSRLCLPMSSFVSGEGR-IILYLYV |
|  | 60 | UniRef90_A0A7K6ZIQ3_start_3_end_496_Evalue_1e-245 | ATLATLKT------RRSHLCLPMSSFASGDGR-VVLYLYV |
|  | 61 | UniRef90_A0A7K7VD55_start_32_end_523_Evalue_1e-255 | ATLATLKS------RRSHLCLPMSSFASSDGR-VVLYLYV |
|  | 62 | UniRef90_A0A444U2D8_start_29_end_532_Evalue_1e-265 | ATLVTLRTYP----WQSKVCLPHFGLISGDGK-IILYLYL |
|  | 63 | UniRef90_A0A6P9CJ49_start_186_end_717_Evalue_1e-284 | ATLATMRT------HKSRLCLPISSLISGEGK-IIPYLYV |
|  | 64 | UniRef90_A0A6J0UY41_start_87_end_623_Evalue_2.3e-287 | ATLATMRT------HKSRLCLPISSLVNREGK-IIPYLYV |
|  | 65 | UniRef90_A0A7K4XA58_start_1_end_361_Evalue_2.5e-167 | ATLVTLRT------RGSRLCLPMASLPSGQGR-LILYLYL |
|  | 66 | UniRef90_H3ACI0_start_28_end_537_Evalue_2.7e-273 | ATLATLRT------WKSRICLPLFSLTGGDGK-IILYLYV |
|  | 67 | UniRef90_A0A7K4LZE5_start_3_end_503_Evalue_2.8e-266 | ATLATLKT------RRSHLCLPMSSFSSGDGR-IVLYLYV |
|  | 68 | UniRef90_A0A851VRD7_start_3_end_426_Evalue_2.9e-158 | ATLATLRT------RGSRLCLPMASPPGGQGK-LIGGGDV |
|  | 69 | UniRef90_A0A7K5GA52_start_25_end_545_Evalue_2.9e-274 | ATLATLKT------RGSRLCLPMSSFASGEGK-IILYLYV |
|  | 70 | UniRef90_A0A8B7J5L7_start_168_end_679_Evalue_2.9e-277 | ATLATLKT------RRSRLCLPMSSFVSSEGR-VILYFYV |
|  | 71 | UniRef90_A0A7E6DXS2_start_95_end_564_Evalue_2.9e-297 | ATLATLKT------SKSHLCLPMSSLVAGEGK-AVLSLYV |
|  | 72 | UniRef90_A0A8D2KZZ1_start_8_end_542_Evalue_2e-274 | ATLATLRT------HNSRLCLPLSSLVSGEGK-IIPYLYV |
|  | 73 | UniRef90_H2M568_start_16_end_529_Evalue_3.1e-217 | AALLTTRVYP----HRDGPCLRHFGLGTGQAR-VVLYVYL |
|  | 74 | UniRef90_A0A6P3VVJ0_start_37_end_570_Evalue_3.4e-213 | AVLVTLRAYP----RSSGPCLPHFGLTGNRAKVVVLYVYL |
|  | 75 | UniRef90_A0A3Q3DYR6_start_15_end_538_Evalue_3.4e-215 | AALLIMRAYP----RRDGPCLPHFGLGGGRAR-LVLYAYL |
|  | 76 | UniRef90_A0A0R4IPI9_start_26_end_553_Evalue_3.4e-231 | AALVTLRAYP----RSSGPCLPHFGLTGNKAK-VVLYVYI |
|  | 77 | UniRef90_UPI0003314560_start_1_end_532_Evalue_3.4e-284 | AALATPRA------SKSHLCLPMSSLMVGDGT-VILSLYV |
|  | 78 | UniRef90_UPI001CF07742_start_34_end_504_Evalue_3.4e-302 | ATLGTLKT------SKSHLCLPMSSLTAGEGK-AIVSLYV |
|  | 79 | UniRef90_A0A3Q0GD38_start_1_end_487_Evalue_3.5e-262 | AALATWRT------RRSRLCLPVSRPAAGEGK-LILYLYV |
|  | 80 | UniRef90_A0A7L3CT33_start_16_end_542_Evalue_3.5e-277 | ATLATLKT------RGSRLCLPMSSFASGEGK-IILYLYV |
|  | 81 | UniRef90_A0A673CNX9_start_13_end_547_Evalue_3.7e-225 | AALLTMRAYP----RRDGPCLPHFGLGGGQAR-VVLYVYL |
|  | 82 | UniRef90_A0A6J2VAV6_start_18_end_552_Evalue_3.7e-233 | AALITMRAYP----RTSGPCLPHFGLTGNTAK-VVLYVYL |
|  | 83 | UniRef90_A0A7L4DFQ1_start_1_end_497_Evalue_3.7e-267 | ATLATLKA------HGSRLCLPMSSFASSEGK-IILYLYI |
|  | 84 | UniRef90_A0A7L0EQY2_start_3_end_504_Evalue_3.7e-269 | ATLATLKT------HGSHFCLPMSGFASGEGK-AILYLYV |
|  | 85 | UniRef90_A0A8K1G3R2_start_14_end_469_Evalue_3.8e-210 | ATLATLRT------RGSRLCLPVAGLPRGQGK-LILYLYV |
|  | 86 | UniRef90_A0A401QAA2_start_28_end_539_Evalue_3.9e-239 | ATLATLRTHP----VRSRVCVPLFGCIDGDGN-VILYLYV |
|  | 87 | UniRef90_UPI0005326EB2_start_82_end_462_Evalue_3e-196 | ATLATLKT------RGSRLCLPMSSFASSEGK-IILYLYV |
|  | 88 | UniRef90_A0A8D0DPE8_start_1_end_543_Evalue_3e-286 | ATLATMRT------QKSRLCLPISSLVSGEGK-VVPYLYV |
|  | 89 | UniRef90_A0A7L4K0J5_start_2_end_497_Evalue_4.1e-248 | AALATLGS------RGSRLCLPVSGFASGRGK-VILYLYV |
|  | 90 | UniRef90_UPI0018642C90_start_55_end_569_Evalue_4.2e-160 | ATLVTLRPRL----RRAPPCLPCAGLVGALAG-PALYLYL |
|  | 91 | UniRef90_A0A1A8I6J6_start_32_end_548_Evalue_4.2e-217 | AALLTMRAYP----RREGPCLPHFGLGGGQAR-AVLYVYL |
|  | 92 | UniRef90_A0A7K8JJZ0_start_3_end_499_Evalue_4.2e-264 | ATLATLKT------HGSRLCLPMSSFASDEGK-VILYLYV |
|  | 93 | UniRef90_H3BGH5_start_24_end_518_Evalue_4.6e-198 | AALSAMTKY-----HRSKICVPLLGLINKDGK-AILYAYV |
|  | 94 | UniRef90_A0A852KNZ0_start_31_end_520_Evalue_4.6e-249 | ATLATMKT------RGSRLCLPMSSFAGGEGK-VILYLYV |
|  | 95 | UniRef90_A0A7K7GGD1_start_23_end_475_Evalue_4.7e-210 | ATLATLRT------RGSRLCLPMASLPGGQGK-LILYLYV |
|  | 96 | UniRef90_A0A7K6VTA4_start_14_end_509_Evalue_4.8e-224 | ATLATMRT------RGSRLCLPMASLPSGQGK-LILYLYV |
|  | 97 | UniRef90_A0A7K6M7M9_start_9_end_497_Evalue_4.9e-219 | ATLATLRA------RGSRLCLPMASLPSGQGR-LILYLYV |
|  | 98 | UniRef90_A0A852B0N9_start_1_end_337_Evalue_5.1e-150 | ATLATLHT------RGSRLCLPMASLPSGQGR-FILYLYV |
|  | 99 | UniRef90_UPI0010A98A13_start_1_end_356_Evalue_5.1e-192 | ----------------SHLCLPMSSFAGGKGK-IIHYLYV |
|  | 100 | UniRef90_A0A6F9BQ47_start_10_end_516_Evalue_5.1e-198 | AALLTLRAYP----TKEGPCLPHFGLTGGRAR-VVLYVYL |
|  | 101 | UniRef90_H2UY98_start_11_end_541_Evalue_5.5e-217 | AALLTMRAYP----HRDGPCLPHFGLRGDQAR-VVLYVYL |
|  | 102 | UniRef90_UPI001CFA05C8_start_25_end_517_Evalue_5.9e-189 | AAFSAMNKY-----HGSQVCVPLIGLINEGGK-AILYIYV |
|  | 103 | UniRef90_UPI00071A4BD4_start_31_end_484_Evalue_5e-215 | ATLATLRT------RGSRLCLPMASLPSGQGK-LILYLYV |
|  | 104 | UniRef90_UPI001CFABF3D_start_34_end_536_Evalue_5e-269 | ATLATLRT------WRSRICLPMFSLINEDGK-IILYLYI |
|  | 105 | UniRef90_A0A851JDQ5_start_1_end_383_Evalue_6.5e-164 | ATLATLRT------RGSRLCLPMASLPKGQGK-LILYLYV |
|  | 106 | UniRef90_A0A811YLR5_start_1_end_485_Evalue_6.5e-181 | AALGALRA------SKAQLCLPLSGLPAAEGA-AAPSLYA |
|  | 107 | UniRef90_A0A7K4JCQ1_start_1_end_505_Evalue_6.6e-266 | ATLATLRS------RSSHLCLPMSSFTGGQGK-VILYLYV |
|  | 108 | UniRef90_A0A091G4V3_start_95_end_447_Evalue_6.7e-180 | -----------------RLCLPVSGFAGGQRK-VILYLYV |
|  | 109 | UniRef90_W5LSM2_start_5_end_553_Evalue_6.9e-229 | AVLVTLRAYP----RTPGPCLPHFGLTGNQAK-VVLYVYL |
|  | 110 | UniRef90_A0A852PRV3_start_14_end_509_Evalue_6e-223 | ATLATLRT------RGSRRCLPMASLASGQGK-VILYLYV |
|  | 111 | UniRef90_A0A7L4GKN4_start_28_end_546_Evalue_7.1e-275 | ATLATLKT------RGSRLCLPMSSFASGEGK-VILYLYV |
|  | 112 | UniRef90_UPI0014027943_start_59_end_478_Evalue_7.4e-88 | AALAALRGHTRGPGPGPQLCLPLLS----GG-LVLYAYA |
|  | 113 | UniRef90_A0A7K6ARY7_start_3_end_509_Evalue_7.7e-270 | ATLATLKT------RSSRLCLPMSSFTSSQGK-IILYLYV |
|  | 114 | UniRef90_A0A401NTY6_start_36_end_533_Evalue_7e-197 | ATLSAMNKY-----HSSRICVPLLGLMNEEGK-AILYTYV |
|  | 115 | UniRef90_A0A7L2RVA3_start_1_end_385_Evalue_8.4e-165 | ATLATLRS------QRSRRCLPMASLASGRGR-LVLYLYL |
|  | 116 | UniRef90_A0A8C5UIR7_start_8_end_472_Evalue_8.6e-216 | ATMATLRT------RGSRLCLPMAGLPSGQGK-LILYLYV |
|  | 117 | UniRef90_A0A556UFX8_start_31_end_546_Evalue_8.9e-213 | AALVTLRAYP----RSVGPCLPHFGLTGNQAK-VVLYVYL |
|  | 118 | UniRef90_A0A3B3T6D4_start_20_end_542_Evalue_9.1e-225 | AALVTLRAHP----WTSRPCLPRLGLVGGEGR-ATLCLYL |
|  | 119 | UniRef90_K7F1D9_start_19_end_550_Evalue_9e-302 | AILATLRT------RKSRLCLPMSSLISGEGK-IILYLYV |

**Figure S5 | Species conservation of GPR75 residues 4.57-5.44.** The multiple sequence alignment was generated using ConSurf (23) using the wt sequence of human GPR75 as query.

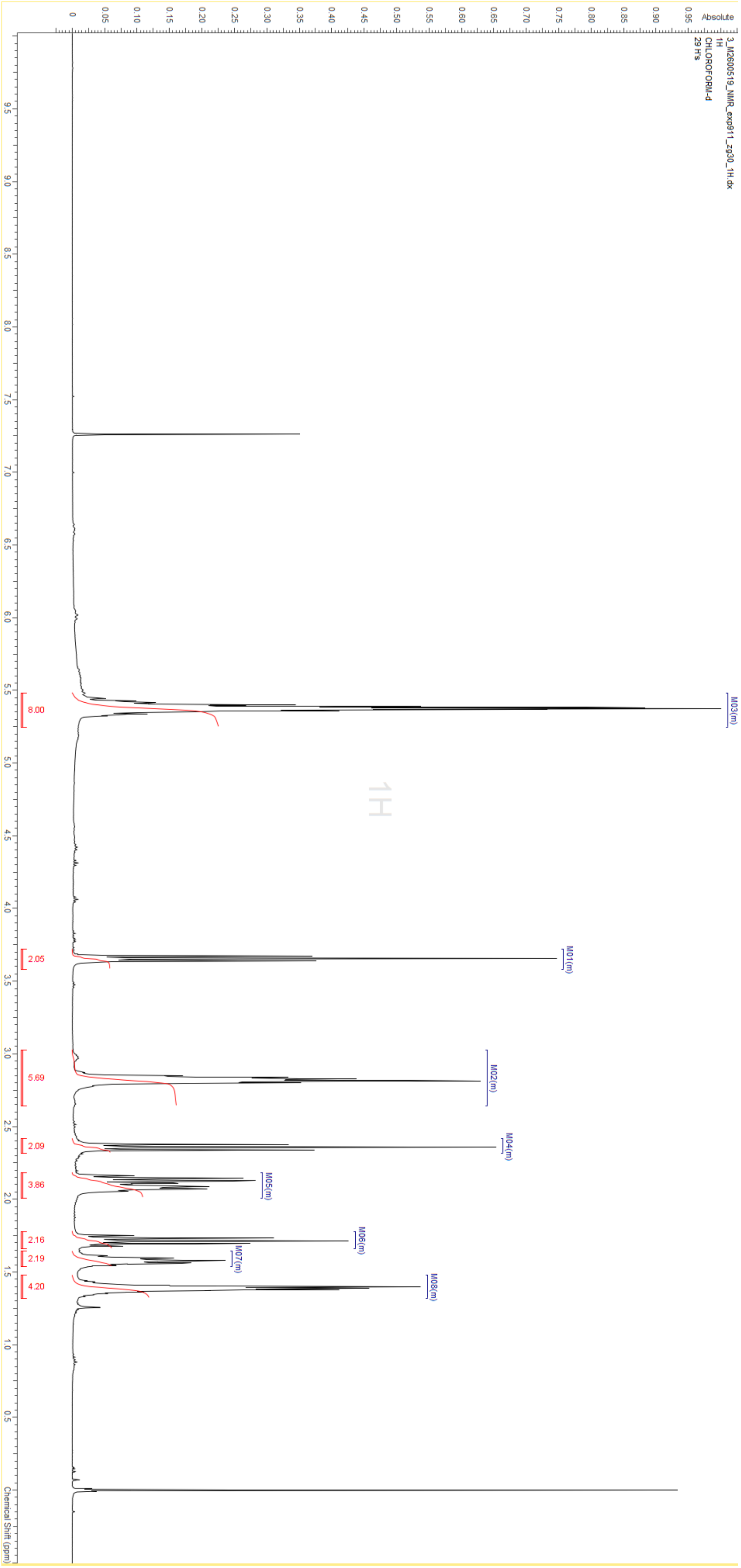

**Figure S6 – 1H-NMR spectrum (400Hz) of 20-HETE.**

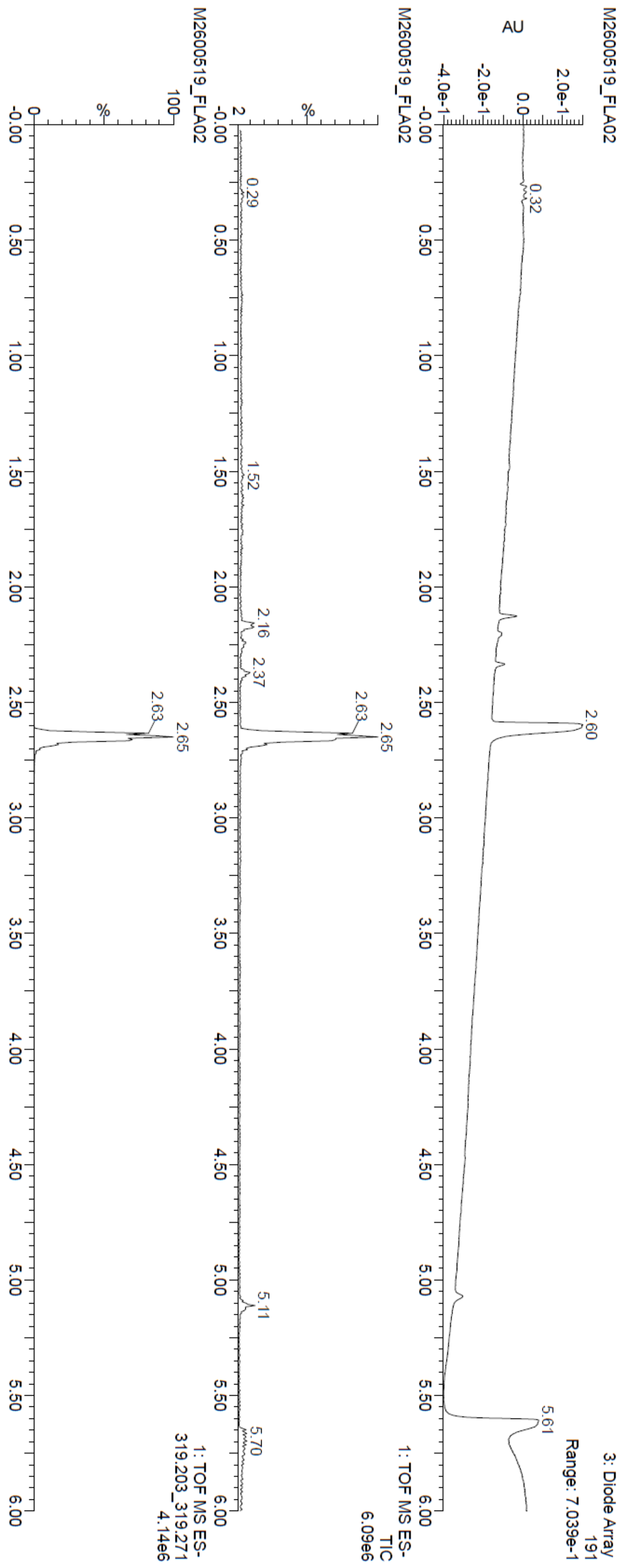

**Figure S7 | HPLC-MS spectra of 20-HETE**

### Supplementary Tables

**Table S1 | Cryo-EM data collection, refinement, and validation statistics**

| | GPR75 wt (PDB 9H2F) |
|---|---|
| **Data collection and processing** | |
| Magnification | 130,000 |
| Voltage (kV) | 300 |
| Electron exposure (e–/Å$^2$) | 40 |
| Defocus range (μm) | -0.5 to -2.0 |
| Pixel size (Å) | 0.951 |
| Symmetry imposed | P1 |
| Initial particle images (no.) | 7,651,212 |
| Final particle images (no.) | 691,411 |
| Map resolution (Å) | 3.43 (focused GPCR map)<br>2.50 (focused map Fab-Nb)<br>2.59 (focused map whole complex)<br>2.65 (consensus map) |
| FSC threshold | 0.143 |
| Map sharpening B factor (Å2) | -90 (focused GPCR map)<br>-98 (focused map Fab-Nb)<br>-93 (focused map whole complex)<br>-109 (consensus map) |
| **Refinement** | |
| Initial model used (PDB code) | none / 7TUY |
| Model composition | |
| Non-hydrogen atoms | 6884 |
| Protein residues | 894 |
| Ligands | 0 |
| *B* factors (Å$^2$) | |
| Protein (min/max/mean) | 1.65/152.85/51.58 |
| R.m.s. deviations | |
| Bond lengths (Å) | 0.007 |
| Bond angles (°) | 1.104 |
| Validation | |
| MolProbity score | 1.53 |
| Clashscore | 5.69 |
| Poor rotamers (%) | 0.39 |
| Ramachandran plot | |
| Favored (%) | 96.55 |
| Allowed (%) | 3.45 |
| Disallowed (%) | 0.00 |

**Table S2 | Resources used and generated in this study**

| REAGENT or RESOURCE | Source | Identifier |
|---|---|---|
| **Biological samples** | | |
| Male rat aortic rings | This study | N/A |
| **Chemicals, peptides, and recombinant proteins** | | |
| LMNG | Anatrace | #NG310 |
| Cholesteryl Hemisuccinate Tris salt | Anatrace | #CH210 |
| cOmplete ULTRA EDTA-free protease inhibitor cocktail | Roche | #06538282001 |
| Dnase I | Sigma Aldrich | #10104159001 |
| 20-HETE | Cayman Chemical | #90030 |
| CCL5 | R&D Systems | #278-RN/CF |
| CCL3 | R&D Systems | #270-LD |
| U-46619 | Sigma Aldrich | #D8174 |
| **Critical commercial assays & consumables** | | |
| Streptactin XT 4Flow High Capacity Resin | IBA Lifesciences | #2-5030-025 |
| Superose 6 increase 10/300 GL | Cytiva | #29-0915-96 |
| R2/1 Au 300 Holey carbon grids | Quantifoil | #N1-C15nAu30-01 |
| FLIPR Calcium 6 assay kit | Avantor | #MLDVR8190 |
| KappaSelect beads | GE Healthcare | #17-5458-01 |
| Protino Ni-NTA agarose resin | Macherey-Nagel | #745400 |
| PD-10 columns | Cytiva | #17085101 |
| NAP-5 columns | Cytiva | #17085302 |
| High sensitivity capillaries | NanoTemper | #PR-C006 |
| Nunc™ MicroWell™ 96 Wells, Nunclon Delta-behandelt | Thermo Fisher Scientific | #136101 |
| 384 well plate | Perkin Elmer | #6007680 |
| PathHunter eXpress CCR5 CHO-K1 ß-Arrestin GPCR Assay (2-Plate) | DiscoverX | #93-0224E2CP0M |
| **Deposited data** | | |
| Composite EM map of GPR75 wt, anti-BRIL-Fab, anti-Fab Nb | This study | EMDB EMD-51800 |
| Coordinates of GPR75 wt, anti-BRIL-Fab, anti-Fab Nb | This study | PDB 9H2F |
| Consensus EM map of GPR75 wt, anti-BRIL-Fab, anti-Fab Nb | This study | EMDB EMD-51796 |
| Focused EM map of GPR75 wt | This study | EMDB EMD-51797 |
| Focused EM map of BRIL, anti-BRIL Fab, anti-Fab Nb | This study | EMDB EMD-51798 |
| Electron microscopy density map of GPR75 wt, anti-BRIL-Fab, anti-Fab Nb | This study | EMDB EMD-51799 |

| | | |
|---|---|---|
| **Cell lines** | | |
| Trichoplusia ni High Five cells | N/A | N/A |
| CHO-3E7 cells | N/A | N/A |
| E.coli BL21 (DE3) cells | N/A | N/A |
| HEK293H cells | N/A | N/A |
| CHO-K1 cells | N/A | N/A |
| CHO-K1 GPR75 cells | This study | N/A |
| ESF cells | Nuvisan GmbH | N/A |
| **Cell culture reagents** | | |
| OptiPRO SFM | Gibco | #12309-050 |
| TransIT Pro | Mirus | #MIR5760 |
| BalanCD Transfectory CHO medium | Irvine Scientific | #91147 |
| GlutatMAX | Invitrogen | #35050-038 |
| Irvine Feed Transfectory Supplement | Irvine Scientific | #91148 |
| Anti-Clumping reagent | Gibco | #0010057DG |
| CHO CD Efficient Feed B | Gibco | #A10240-01 |
| Sartoclear Dynamics Lab Filter Aid | Sartorius | #SDLV-0500-20C—E |
| DMEM high glucose, pyruvate | Gibco | #11995-073 |
| Pen/Strep | Gibco | #15149-122 |
| 100x L-glutamine 200 mM | Gibco | #25030-024 |
| Heat inactivated FBS | Gibco | #10082-147 |
| 1x Trypsin-EDTA 0.05% | Gibco | #25300-062 |
| DPBS | Gibco | #14190-250 |
| OptiMEM | Gibco | #31985-062 |
| Lipofectamine 2000 | Invitrogen | #11668019 |
| Versene | Gibco | #15040-066 |
| HBSS 10x | Gibco | #14065-056 |
| HEPES 1M | Gibco | #15630-080 |
| Prolume Purple | NanoLight | #369-10 |
| Non-enzymatic cell dissociation solution | Sigma Aldrich | #C5789 |
| Ham’s F-12 (Kaighn’s) growth medium | Gibco | #21-127-022 |
| Forskolin | Abcam | #AB1200585MG |
| cAMP assay kit | Perkin Elmer | #502118313 |
| Geneticin | N/A | N/A |
| Zeocin | N/A | N/A |
| FuGene | Promega | #E2311 |
| **Recombinant DNA** | | |
| Plasmid: pFastBac1 GPR75 wt | This study | N/A |
| Plasmid: pFastBac1 GPR75 C189S | This study | N/A |
| Plasmid: pFastBac1 GPR75 L190A | This study | N/A |
| Plasmid: pFastBac1 GPR75 P191A | This study | N/A |
| Plasmid: pFastBac1 GPR75 M192A | This study | N/A |
| Plasmid: pFastBac1 GPR75 AAAA | This study | N/A |
| Plasmid: pTT5 anti-BRIL Fab heavy chain | This study | Mukherjee et al. |

| | | |
|---|---|---|
| Plasmid: pTT5 anti-BRIL Fab light chain | This study | Mukherjee et al. |
| Plasmid: pET24 anti-Fab Nb | This study | Ereño-Orbea et al. |
| Plasmid: pcDNA3.1+ GPR75 wt | This study | N/A |
| Plasmid: pcDNA3.1+ GPR75 C189S | This study | N/A |
| Plasmid: pcDNA3.1+ GPR75 L190A | This study | N/A |
| Plasmid: pcDNA3.1+ GPR75 P191A | This study | N/A |
| Plasmid: pcDNA3.1+ GPR75 M192A | This study | N/A |
| Plasmid: pcDNA3.1+ GPR75 AAAA | This study | N/A |
| Plasmid: pcDNA3.1+ GPR20 wt | This study | N/A |
| Plasmid: pcDNA3.1+ GPR52 wt | This study | N/A |
| Plasmid: pcDNA3.1+ S1P3 wt | This study | N/A |
| Plasmid: pcDNA3.1+ TBXA2R wt | This study | N/A |
| Plasmid: pcDNA3.1+ Gai1-RLuc8 | This study | Olsen et al. |
| Plasmid: pcDNA3.1+ Gaq-RLuc8 | This study | Olsen et al. |
| Plasmid: pcDNA3.1+ Gß3 | This study | Olsen et al. |
| Plasmid: pcDNA3.1+ Gg9-GFP2 | This study | Olsen et al. |
| **Software and algorithms** | | |
| Unicorn 7.8.0 | Cytiva | #29702890 |
| EPU version 3 | Thermo Fisher Scientific | N/A |
| cryoSPARC version 4.5.3 | Structura Biotechnology | N/A |
| ChimeraX version 1.6.1 | UCSF | N/A |
| PHENIX version 1.21.1 | Afonine et al., 2018 | https://phenix-online.org/ |
| ColabFold | Mirdita et al., 2022 | N/A |
| PR.ThermControl version 2.1.1 | NanoTemper | N/A |
| GraphPad Prism version 10.1.2 | GraphPad Software | https://www.graphpad.com |
| IOX version 2.9.5 | N/A | N/A |
| **Instruments** | | |
| Akta Pure 25M | Cytiva | #29018226 |
| PELCO EasyGlow Glow Discharge Cleaning System | Ted Pella | 91000 |
| Vitrobot Mark IV | Thermo Fisher Scientific | N/A |
| Glacios 200 kV transmission electron microscope | Thermo Fisher Scientific | N/A |
| Titan Krios G4 300 kV transmission electron Microscope, Selectris energy filter, Falcon 4i camera, IMP Vienna | Thermo Fisher Scientific | N/A |
| Prometheus NT.48 | NanoTemper | N/A |
| Dionex IFC-UltiMate-3000 | Thermo Fisher Scientific | N/A |
| Heracell 240i CO2 incubator | Thermo Fisher Scientific | #51032875 |
| BioTek ELx405 select microplate washer 96-well | Agilent | N/A |
| MultiDrop Combi | Thermo Fisher Scientific | #5840330 |
| PheraStar FSX incl. BRET2 Plus module | BMG Labtech | N/A |
| emkaBATH4 | EMKA Technologies | N/A |
| it-1 force transducer | EMKA Technologies | N/A |

318
